# Collective Microbial Effects Drive Toxin Bioremediation and Enable Rational Design

**DOI:** 10.1101/2025.03.28.645802

**Authors:** Mahmoud Yousef, Kiseok Keith Lee, Jonathan Tang, Vasileios Charisopoulos, Rebecca Willett, Seppe Kuehn

## Abstract

The metabolic activity of microbial communities is essential for host and environmental health, influencing processes from immune regulation to bioremediation. Given this importance, the rational design of microbiomes with targeted functional properties is an important objective. Designing microbial consortia with targeted functions is challenging due to complex community interactions and environmental heterogeneity. Community-function landscapes address this challenge by statistically inferring impacts of species presence or absence on function. Similar to fitness landscapes, community-function landscapes are shaped by both additive effects and interactions (epistasis) among species that influence function. Here, we apply the community-function landscape approach to design synthetic microbial consortia to degrade the toxic environmental contaminant bisphenol-A (BPA). Using synthetic communities of BPA-degrading isolates, we map community-function landscapes across increasing BPA concentrations, where higher BPA means greater toxicity. As toxicity increases, so does epistasis, indicating that collective effects become more important in degradation. Further, we leverage landscapes to rationally design communities with predictable BPA degradation dynamics *in vitro*. Remarkably, designed synthetic communities are able to remediate BPA in contaminated soils. Our results demonstrate that toxicity can drive epistatic interactions in community-function landscapes and that these landscapes can guide microbial consortia design for bioremediation.

## Introduction

Microbial communities drive processes essential for human and environmental health. Humans have manipulated environmental conditions for millennia to recruit microbial communities with desirable properties, from food fermentation [1] to wastewater treatment [2]. However, using wild or enriched communities to perform a functional role of interest faces challenges. For example, fecal microbial transplants can effectively treat deadly infections [3, 4], but patient outcomes can be variable and sometimes detrimental [5]. Similarly, in a remediation context, where enriched communities can decontaminate pollutants in soils or water, the process is often slow [6, 7] and challenging for decontamination of complex or toxic compounds such as polycyclic aromatic hydrocarbons [8, 9].

These challenges have driven recent interest in synthetic ecology: engineering communities with desirable functional properties. However, rationally designing such consortia is difficult because community function is affected by a myriad of factors, including gene expression [10], ecological interactions [11–13], and environmental variables [14]. As a result, dissecting the relationship between the species present, their abundances, and higher-level community community functions such as metabolism can be a challenge.

One route to overcoming this challenge is to exploit a conceptual analogy with fitness landscapes. Community-function landscapes, which map community composition to function [13, 15, 16], borrow directly from the notion of fitness landscapes in genetics which map a genotype to an organism’s fitness [17]. Remarkably, this simple approach can accurately predict how community function depends on composition [15] through elucidating additive effects (average effect of a strain on a community’s function) and epistatic interactions (average effect of a pair or group of strains on a community’s function beyond their individual additive effects) [18].

However, with few exceptions [19], most work inferring community-function landscapes targets simple synthetic communities [20] and does not solve critical applied problems in biotechnology. Furthermore, community-function landscapes are typically inferred in a single environmental condition [15, 19]. It is unclear how the landscape structure depends on fluctuating environmental conditions such as nutrient availability. The dependence of the landscape on the environment is especially critical for the deployment of engineered communities in natural environments where resident consortia and variable environmental conditions might render any designed consortia ineffective [21]. Thus, there remains a pressing need to test the approach on a challenging applied community design problem where environmental variation might play a role.

One biotechnological process that could benefit from environment-influenced rational community design is bioremediation. In this context, dangerous chemicals that contaminate water and soil environments can be remediated by the metabolic activity of microbes [22]. Contaminant removal is commonly approached through both single-strain and community-level bioremediation [6, 23 – 25]. However, communal bioremediation attempts are limited due to a lack of rational design principles to engineer communities [14, 26, 27].

One major target of bioremediation efforts is bisphenol-A (BPA), a widely used industrial chemical and pollutant [28, 29] whose removal offers significant health benefits [30]. Bacteria can utilize BPA as a carbon source during aerobic respiration[31–33]. Although BPA degradation by individual bacterial strains has been extensively studied [34, 35], many fail to achieve complete degradation [34] due to the toxicity of BPA and the byproducts of BPA degradation [36–40]. Some studies suggest that communities can overcome the toxicity of BPA and its intermediates [41], however, there is no rational route to selecting strains to include in BPA degrading consortia. Part of the challenge of selecting strains is that we lack definitive knowledge of the genes responsible for every step of a BPA degradation pathway, as only a handful of steps have been characterized [31–33, 38, 42, 43]. Furthermore, there has been little study of the role of environmental factors in BPA bioremediation [34].

Here we address these challenges by exploiting a community-function landscape approach to design BPA-degrading communities across a range of increasing BPA concentrations. Inferred landscapes show that collective effects, which arise in landscapes through epistatic statistical interactions between taxa, increase with higher levels of the contaminant. In addition, we find that the structure of the landscape changes smoothly as the BPA concentration rises. We then use this landscape to design communities with desired rates of BPA degradation *in vitro*. Finally, we show that our designed communities remediate BPA in contaminated soils. Together, our results highlight the power of community function landscapes in understanding community function under varying environmental conditions and as a tool for the statistical engineering of communities with real-world applications.

## Results

### A library of isolates to enable community design for remediation

We set out to design synthetic communities of bacteria capable of degrading BPA. First, we assembled communities from a library of bacterial soil isolates available in our laboratory [44] (largely proteobacteria) and tested their ability to degrade BPA in minimal media. BPA degradation was assessed by sampling cultures in time and using a colorimetric assay to measure BPA concentration in samples (see Methods). We found that none of these communities could degrade BPA (Fig. S2), indicating that the ability to degrade BPA is not a common trait for readily cultured soil isolates.

Therefore, we sought to construct a strain bank of bacterial isolates with the capacity for BPA degradation. Our goal was to acquire 10-20 strains that could be used to assemble synthetic communities. To accomplish this, we identified five sites with potential BPA contamination—two sites in proximity to BPA-producing factories and three sites downstream of these factories (for details of the sampling sites and methods see Methods and Table S2, rows 1-5).

We enriched each of the five soil samples for BPA degradation by aerobically incubating 3 grams of soil in 60 mL M9 media with 30 ppm BPA as the sole carbon source (see Methods). All of the samples exhibited BPA degradation and continued to do so upon serial dilutions. From these five samples, we isolated and identified 16 bacterial strains, ten of which degrade BPA from 30 ppm in monoculture (degraders) and 6 that do not degrade on their own (non-degraders) but were retained due to being co-isolated with BPA degraders (Fig. 1B, Table S3). We confirmed BPA degradation occurs via aerobic respiration by measuring CO_2_ production in monocultures of each of our ten degrading isolates (Fig. S3).

**Figure 1:**
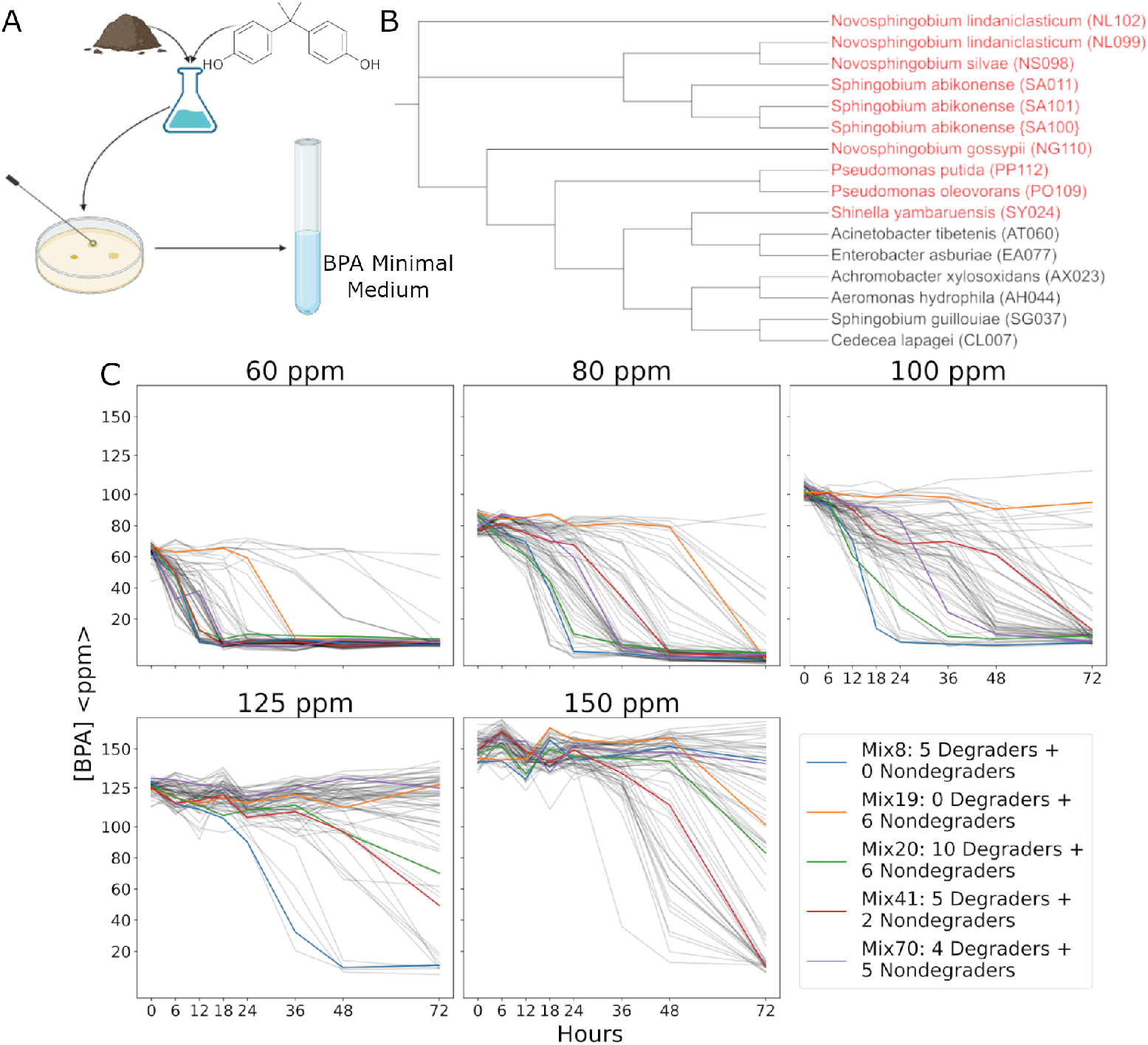
Isolates enable construction of synthetic bacterial communities that degrade BPA across varying initial concentrations. **(A)** Protocol for enrichment of soil samples. After enrichment, isolates are streaked to purity, and then screened for BPA degradation. **(B)** Phylogenetic tree of the 16 strains isolated. Red strains denote strains that degrade BPA on their own from an initial concentration of 30 ppm. **(C)** BPA degradation curves of 70 communities in five initial concentrations of BPA. Five communities are highlighted: Mix8 (blue), Mix19 (yellow), Mix20 (green), Mix41 (red), and Mix70 (purple). All measurements are corrected for evaporation (Methods).

### Structure of community-level degradation with increasing toxicity

Armed with a library of isolates capable of BPA degradation we next set out to assemble synthetic communities and test their BPA degradation capabilities. Previous studies [19] suggest that randomly assembling diverse communities and quantifying their functional property of interest can enable statistical design. This approach works especially well when community-function landscapes are not rugged, and predicting community function requires only estimating the average effect of each strain on function [15].

Therefore, we assembled 70 communities from our 16 isolates and inoculated them into BPA minimal medium. These communities had a mean richness of 6 (with a standard deviation of 3) and included a community of all 16 strains, a community of all 10 degraders, and a community of all 6 non-degraders. The remaining 67 communities were assembled randomly. To assemble communities, each bacterial isolate was grown axenically, and then the appropriate strains were combined with a fixed initial OD of 0.02 for each strain irrespective of community richness.

For each community, we evaluated its capacity to degrade BPA by incubating it for 72 hours in a BPA minimal medium and monitoring BPA concentrations over time (see Methods). Previous work suggests that increasing BPA concentration is toxic to bacteria[36, 37], so we expect that increasing the initial concentration renders degradation more challenging. Therefore, we measured every community’s BPA degradation dynamics at five initial BPA concentrations (60, 80, 100, 125, and 150 ppm). Note that while typical environmental concentrations are often below 1 ppm [29], contaminated BPA sites often range in BPA concentrations between approximates 10 and 200 ppm [45]

Figure 1 C shows the degradation dynamics for all 70 communities in five initial BPA concentrations. Five representative communities are highlighted in color across all concentrations. From these data, we made several initial observations. First, the community consisting of the six non-degrading isolates (Mix19) degrades BPA from an initial concentration of 60 ppm, suggesting the role of collective effects in degradation. Second, as initial BPA concentration increases, it becomes increasingly harder for our communities to degrade. For example, note that most communities do not degrade BPA when incubated at 150 ppm (Fig 1 C, bottom-center). Third, we observe that many communities that were able to degrade BPA at low concentrations cannot do so at higher concentrations (Mix19 and Mix70, orange and green traces, Figure 1 C).

To quantify variation in community BPA degradation dynamics, we computed the area under the curve (AUC) for the BPA concentration in time. Communities that degrade BPA rapidly have a low AUC and the converse (Fig. 2 A). We chose this metric over explicit modeling of BPA degradation dynamics due to the complexity of the BPA dynamics we observed (such as differing timescales of BPA degradation and plateaus mid-way through degradation).

**Figure 2:**
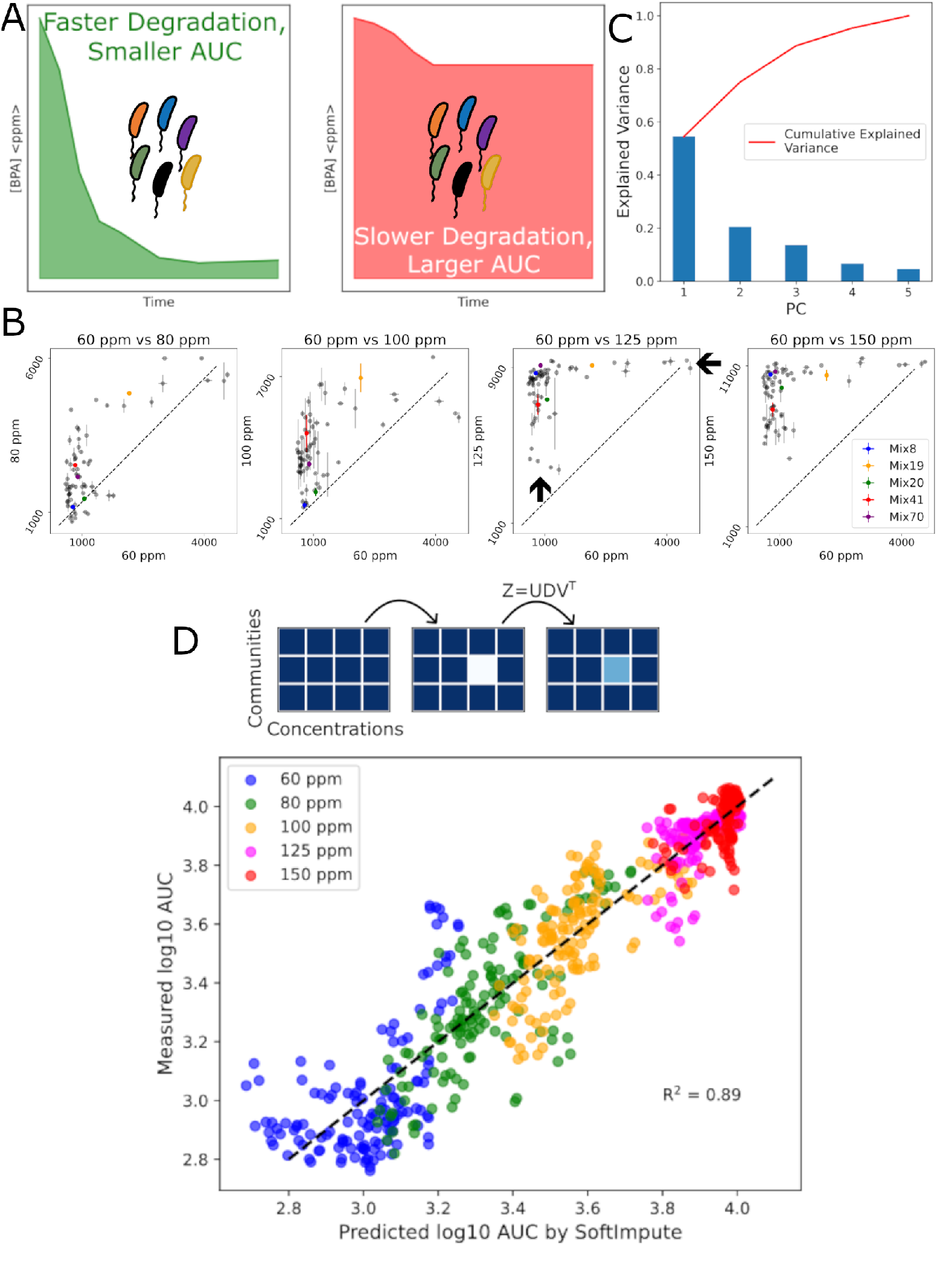
BPA Degradation Varies in a Low Dimensional Manner Across Concentrations. **(A)** Area Under the Curve (AUC) allows us to quantify BPA degradation of communities. A smaller AUC indicates faster and more complete degradation (left), while a higher AUC indicates the opposite (right). **(B)** Comparing the AUCs of our 70 communities between 60 ppm initial BPA and each of the other four initial concentrations (80, 100, 125, 150 ppm). Vertical error bars represent the standad deviation of AUC across technical replicates. Dotted lines indicate normalized equivalent AUC. Two modes of variation become clearer as the difference in concentrations increases (arrows, bottom left). **(C)** PCA analysis shows that the first two Principal Components explain 79% of the variation in communities’ AUCs across concentrations. **(D)** (TOP) Schematic of SoftImpute predictions for AUC. A matrix where each row represents a community and each column an initial BPA concentration with entries being BPA degradation AUC. A single entry is deleted from the matrix, and SVD is used to predict the deleted entry (see Methods). (BOTTOM) Predictions of all AUCs for all communities in all 5 initial concentrations. Predictions are performed with a *log*_10_ transformation of AUC. Points are colored by the initial concentration. R^2^ = 0.89.

The AUC metric enables us to directly compare the performance for each community between a low initial BPA concentration (60 ppm) and higher, more stressful concentrations. (Fig. S4 B). Remarkably, every community that did not degrade BPA from 60 ppm also did not degrade at any of the higher concentrations. We also do not observe any community that degrades BPA faster as concentration increases (points below the dashed lines, Fig. 2 B). This shows us that a community’s ability to degrade BPA from a lower initial concentration is necessary but not sufficient for that community to degrade BPA from a higher initial concentration. More broadly, we observe a relatively simple structure of variation in community function as the BPA concentration increases, consisting of two axes (Fig. 2 B 60 vs 125ppm, black arrows). One axis comprises communities that are fast at the lower concentration but spread across AUC values at the higher concentration, and a second axis of communities that are poor degraders at the higher concentration but exhibit variable AUC at the lower concentration. These patterns are also observed when comparing higher concentrations of BPA (Fig. S4). The relatively simple structure in the variation of AUC with BPA concentration suggested a deeper statistical interrogation of how community degradation changed with initial concentration.

### Low-dimensional variation in community function

To further investigate the structure of variation in community function across initial BPA concentrations, we constructed a matrix with rows defined by community composition and columns by initial BPA concentration. The entries of this matrix were the AUC for each community in each BPA concentration. We performed Principal Component Analysis (PCA) on this matrix, which revealed low-dimensional structure in community performance across BPA concentrations with just two dimensions capturing nearly 80% of the variation in the data (Fig. 2C). This result suggests that the changes in community function (AUC) across BPA concentration are low dimensional, akin to soft modes observed widely in biological systems [46].

The observations led us to ask whether the low dimensionality of community AUCs can be exploited to make statistical inferences. In particular, the results in Fig. 2C suggest that low-dimensional variation in the data might be exploited via matrix imputation methods that rely on low-rank approximations[47]. To test this, we took the matrix of our BPA AUCs across the five concentrations for each community and removed a single entry corresponding to an AUC for a single community composition at a single initial BPA concentration. We then imputed (predicted) the missing value via matrix imputation that exploits a low-rank approximation to matrices with missing data (`SoftImpute` [47]). We then repeated this iteratively for each community at each concentration (Fig. 2D, see Methods). Fig. 2 D shows the measured versus predicted AUC using the `SoftImpute`. These imputed values are very close to the measured AUC values (*R*^2^ = 0.89, *p*-value < 0.01 as calculated by the `scikit-learn` function `r2 score`). The result shows that the BPA degradation performance of a given community at a given initial BPA concentration can be predicted from knowledge of how that community performs at other BPA concentrations. We also show that this method works when up to 10% of the communities are held out for prediction (Fig. S6). These results suggest a simple structure of the underlying community-function landscape.

### Learning the community-function landscape for BPA degradation

We have shown that variation in BPA degradation is low-dimensional. The imputation method shown in Fig. 2 enables predictions of the performance of a community across BPA concentrations so long as any given community has been measured at least one BPA concentration. However, the imputation approach cannot reveal the roles of individual strains within a community in driving BPA degradation. Nor can imputation be used to rationally design communities with a composition that has not already been measured.

To assess the impact of each species on the degradation of BPA, we inferred community function landscapes, as discussed above. Recent work [15, 16] uses linear regression to measure the impact of strain presence and absence on community function in a manner that is identical to inferring the impact of mutations on phenotypes [13, 17]. Here, we utilized the method developed in Skwara *et al*. [15]. These linear models predict *log*(*AUC*) as a metric of community BPA degradation from strain presence/absence (*x*_*i*_, where *i* indexes species) and include additive terms that capture the average effect of individual strains on degradation (*β*_*i*_) and epistatic terms that capture the (statistical) effects of species-species interactions on *log*(*AUC*) (*γ*_*i,j*_, Fig. 3A, Eqn. 1). Note that because lower AUC corresponds to faster degradation, more negative terms (*β*_*i*_, *γ*_*i,j*_ < 0) indicate better BPA degradation. We fit the coefficients *β*_*i*_ and *γ*_*i,j*_ independently for each of the 5 initial BPA concentrations. Since there are 136 parameters and only ∼70 data points, we employ a ridge regularization during inference, and the optimal regularization hyperparameter was determined by leave-one-out cross-validation. Model predictions were assessed via leave-one-out cross-validation (see Methods, Fig. S5 A). We find that third-order terms do not improve model fits (Fig. S5C).

**Figure 3:**
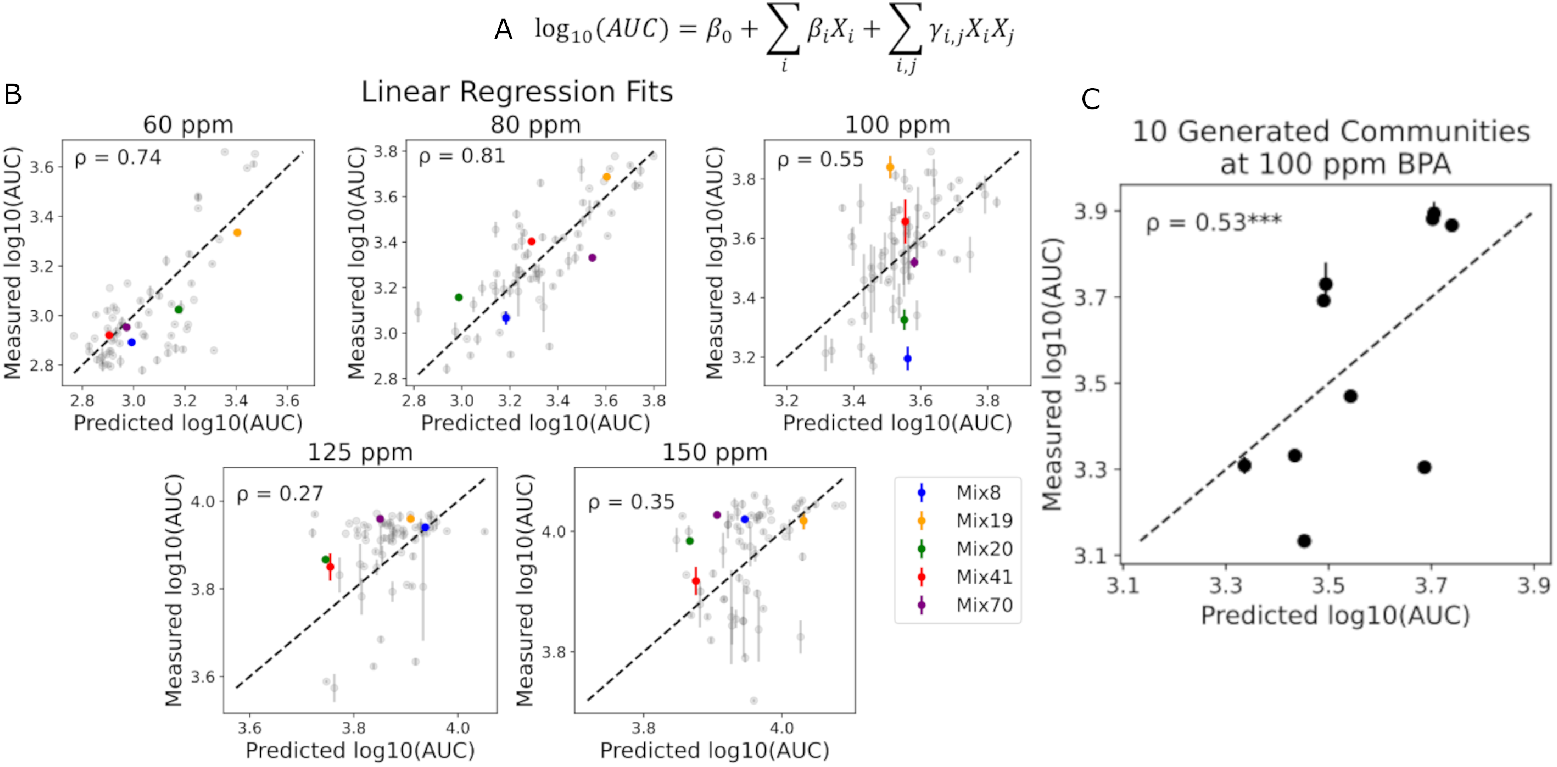
Inferring community-function landscapes enables design. **(A)** The linear regression model for inferring community-function landscapes via regression. The response variable is *log*_10_(*AUC*). The *β*_*i*_ capture additive effects of each strain on function, and the *γ*_*i,j*_ are epistatic contributions. To avoid overfitting, the model is regularized via a Ridge penalty term (see Methods). Independent regressions are performed for each initial BPA concentration. See Methods for details. **(B)** For each concentration, a second-order model of the form shown in (A) is inferred. Hyperparameter tuning is performed via cross-validation. Each panel plots measured versus predicted *log*_10_(*AUC*) for each community at a single initial BPA concentration. For each community, at each concentration, the model is trained on the remaining 69 communities (leave-one-out), and this model is used to predict the *log*_10_(*AUC*) for the held-out community. Both duplicates for each community composition are held out at once. This predicted *log*_*1*0_(*AUC*) is plotted on the axis and the measured *log*_10_(*AUC*) on the y-axis. The dotted lines denote perfect predictions. Vertical error bars represent the standard deviation of *log*_10_(*AUC*) across technical replicates. **(C)** For the model trained in the third panel of the top row of (A), we computationally designed 10 new communities from the model and measured their BPA degradation. Scatter plot of predicted vs measured *log*_10_ *AUC* for these 10 communities at the initial concentration of 100 ppm. Vertical error bars represent the standard deviation of *log*_10_(*AUC*) across technical replicates. *ρ* = 0.53, ***p < 0.001 from 1000 bootstrapping iterations rejects the null hypothesis that the Pearson’s correlation coefficient is ≤0.

The leave-one-out predictions are shown in Fig 3B, where we observe generally good out-of-sample predictions (*ρ* between 0.27 and 0.81). However, we do observe a decrease in the quality of model predictions as initial BPA concentration increases (Fig. 3B). The optimization of these models yields a set of fitted parameters (*β*_*i*_, *γ*_*i,j*_) at each concentration. Collectively, these parameters define the community-function landscape for BPA degradation by this library of strains.

Unlike the imputation methods, community-function landscapes enable us to computationally design new communities from the 16 isolates with predicted high or low *log*10(*AUC*). Using the model fit to 100ppm, we computationally sampled a range of previously unmeasured communities with a range of predicted performance (AUC). We then constructed these communities and measured their BPA degradation dynamics at the appropriate initial BPA concentrations. Remarkably, we find the simple model provides good predictive power of the performance of entirely new communities (Fig 3C, S9). These results establish the fidelity of our statistical approach for predicting BPA degradation as a function of community composition.

### Community-function landscapes reveal that toxicity drives epistasis

One way to characterize community-function landscapes is by the degree to which they are dominated by additive versus epistatic effects. Landscapes with high epistasis (many significant *γ*_*i,j*_ relative to *β*_*i*_) are inherently rugged, with many local minima and maxima. Conversely, purely additive landscapes (*γ*_*i,j*_ = 0)) are smooth and take on a Mount Fuji structure with a single optimum. Previous studies have shown that epistasis is generally weak in synthetic communities [15], leaving open the question of what drives epistasis in community-function landscapes. With this picture in mind, we compared the regression coefficients across initial BPA concentrations (Fig. S7A).

The first property of these landscapes we note is that the additive terms (*β*_*i*_) decline in magnitude as BPA concentration rises. Specifically, at 60 ppm, many of these terms are large and negative, while at concentrations >100 ppm additive coefficients become closer to zero or positive (Fig. 4A, with the exception discussed below). What this means is that the average effects of individual strains on BPA degradation are becoming less important in a statistical sense as BPA concentrations rise. This observation is the first hint that the ruggedness of the landscape changes with BPA concentration.

**Figure 4:**
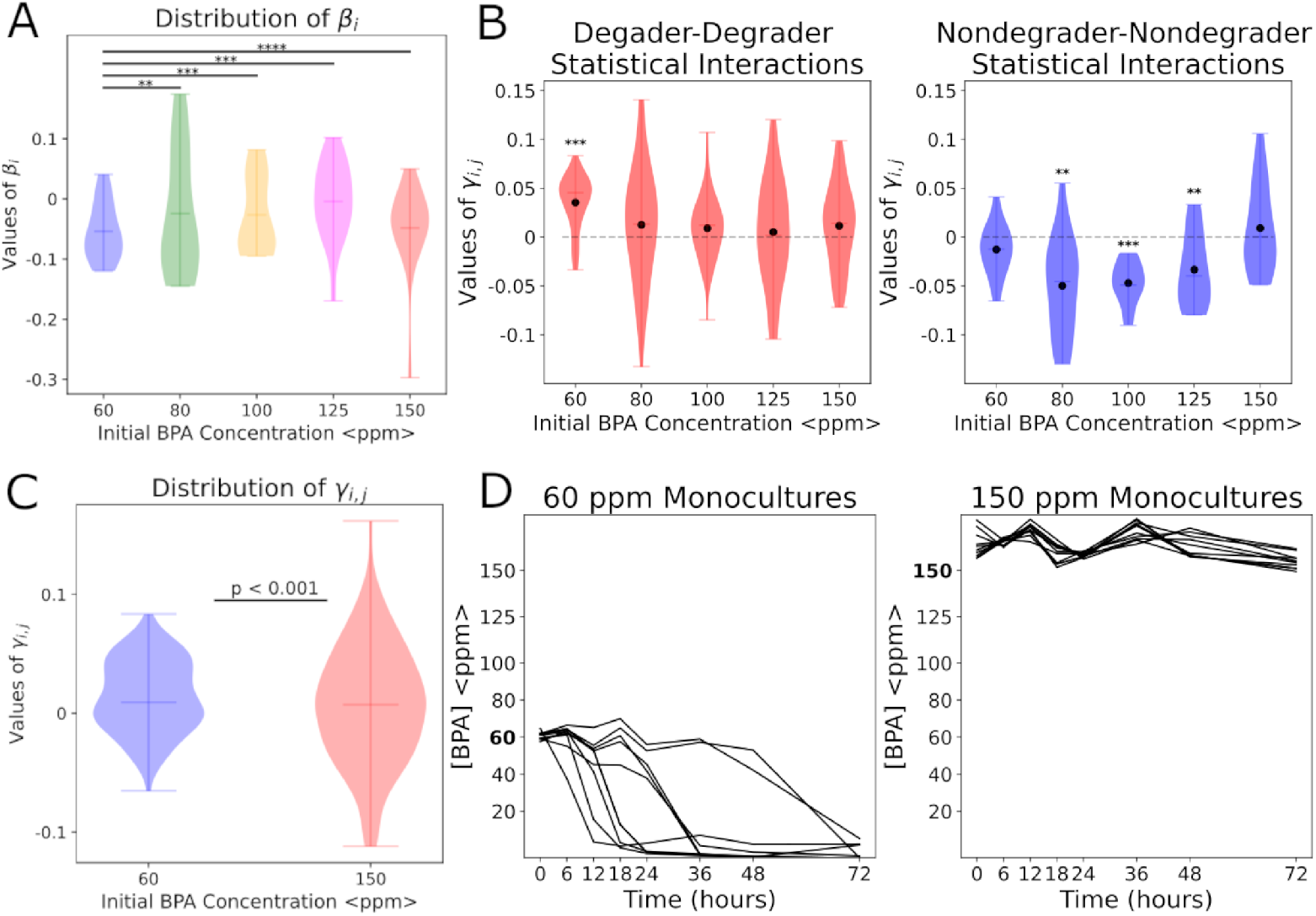
Epistasis Increases as Toxicity Increases. **(A)** Distribution of additive coefficients (*β*_i_) for linear models fit to each of five initial BPA concentrations (Fig. 3). Mean of coefficients rises towards zero as concentration increases. Horizontal lines within the distributions represent the mean of coefficients. **p < 0.01, ***p < 0.001, ****p < 0.001 when strain PO109 is excluded, all determined by a t-test comparing the distribution of additive coefficients at 60 ppm to the additive coefficients of the other four concentrations. **(B)** Breakdown of epistatic coefficients (*γ*_i,j_) by degrader-degrader coefficients (left) and nondegrader-nondegrader coefficients (right). Degrader-degrader terms are competitive (positive, slowing down degradation) at 60 ppm, but become neutral on average as toxicity increases. Nondegrader-nondegrader effective interactions are cooperative (negative, promoting degradation) on average for the intermediate concentrations but neutral at 60 and 150 ppm. Solid dots represent the mean of coefficients. **p < 0.01, ***p < 0.001 from 1000 bootstrapping iterations rejects the null hypothesis that the mean epistatic coefficient is 0. **(C)** Distribution of all epistatic coefficients for models inferred at 60 and 150 ppm. Epistatic coefficients are more dispersed at 150 ppm, indicating the greater presence of epistasis at high BPA toxicity. Horizontal lines within the distributions represent the mean of coefficients. p < 0.001 determined via an f-test. **(D)** Monocultures of the ten BPA-degrading strains in an initial BPA concentration of 60 ppm (left) and 150 ppm (right). None of the monocultures degraded BPA at 150 ppm, indicating the necessity of community effects.

Interestingly, the additive coefficients at 150 ppm show a singular large negative coefficient, which corresponds to strain PO109 (Fig. 4 A, 150 ppm and Fig. S7). This strongly negative coefficient implies that PO109 on average contributes strongly to BPA degradation at a highly toxic concentration whenever it is present in a community; however, we note that PO109 is unable to degrade 150 ppm BPA on its own (Fig. 4 D). Thus, we understand this phenomenon as PO109 strongly contributing to BPA degradation from 150 ppm with high effective interactions with all other strains on average; therefore, this additive term actually captures an overall epistatic effect regarding PO109. We also note that the additive coefficient for PO109 is also strongly negative at 125 ppm (but not as negative as 150 ppm) (Fig. S7 A). This further supports PO109’s supporting role in degrading toxic concentrations of BPA.

Next, we plotted epistatic coefficients representing degrader-degrader and non-degrader-non-degrader statistical interactions across the range of BPA concentrations (Fig. 4B). Epistatic effects (*γ*_*i,j*_) between degrading strains are positive at 60 ppm initial BPA and zero on average for the higher concentrations. This means that at 60 ppm, degraders tend to inhibit each other in terms of BPA degradation, but at higher concentrations of BPA, this is not the case. Moreover, the spread in epistatic terms increases between degraders as the concentration rises. The increasing variance in the distribution of epistatic terms means that there are more extremely negative and positive *γ*_*i,j*_ as BPA concentration rises, indicating stronger epistasis. This is further shown by the increased spread of epistatic terms at 150 ppm compared to epistatic terms at 60 ppm (Fig 4C)

For nondegrader-nondegrader coefficients (Fig. 4B, right panel), we observe a decline in the mean epistatic interaction between non-degrading strains as the concentration rises. This decline means that there is more synergistic epistasis between strains. We note that the trend is true up to 125ppm, but at 150ppm, this epistasis rises (Fig 4 C).

The increased epistasis can be quantified by the F_1_ statistic 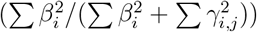 (Fig. 5, top), a metric that quantifies the relative contribution of additive versus epistatic terms to the landscape. Note that if *γ*_*i,j*_ = 0 for all *i, j* then *F*_1_ = 1, but as 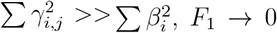. Computing *F*_1_ statistics shows us that epistasis is consistent between 60 and 100 ppm BPA and drops considerably at 125 ppm, showing that epistatic terms dominate as the concentration increases (Fig 5, bottom). The increase in *F*_1_ at 150 ppm is solely explained by the large *β*_*i*_ term corresponding to PO109 (which we discussed above). Removing the contribution from this strain reveals increasing epistasis at 150 ppm.

**Figure 5:**
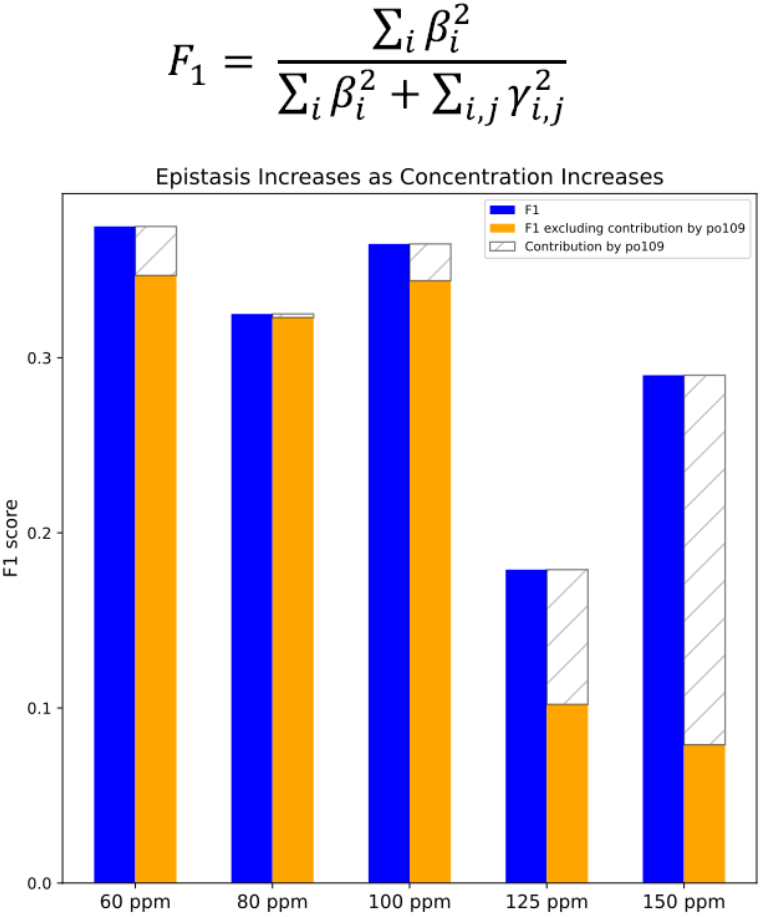
F_1_ Decreasing Shows Increasing Epistasis Driven by BPA Toxicity. **(Top)** Equation for the F_1_ statistic. A higher F_1_ statistic indicates a greater contribution by additive (*β*_i_) terms, thus less epistasis. **(Bottom)** The calculated F_1_ statistic for each of the five models (blue). On average, F_1_ decreases as concentration increases, showing rising epistasis as toxicity increases. Although the F_1_ statistic increases at 150 ppm, this increase is due to the contribution of PO109 alone, and accounting for it maintains the pattern of decreased epistasis (orange and striped)

The conclusion that epistasis rises with BPA concentration strongly suggests that BPA degradation becomes more of a collective, community-level process as toxicity increases. To test this hypothesis, we measured the ability of ten of our strains to degrade BPA in monoculture at 60 ppm and 150 ppm. In agreement with our landscape inferences, we found that while individual strains could degrade BPA at 60 ppm, no single-isolate could degrade at 150 ppm, showing that communities perform better as concentrations rise (Fig. 4D).

Finally, the observation that epistasis increases with concentration and additive coefficients become less important, raises a question regarding the success of the imputation methods shown in Fig. 2 D. In particular, the success of imputation (Fig. 2 D) means that the structure of variation in AUC across BPA concentrations is low-dimensional. It is unclear how this low dimensionality manifests in our inferred community-function landscapes because models for each concentration were inferred independently. Thus, we suspected that the variation in community-function landscapes, changes in *β*_*i*_ and *γ*_*i,j*_ with concentration, should also be low-dimensional. To test this idea, we performed PCA on a matrix of regression coefficients (Methods) and found that 75% variance in coefficients is explained by the first two components (Fig. S7). The result suggests that the community function landscape varies in a simple manner as the concentration of BPA rises.

The low-dimensional variation in landscape coefficients suggests that we might be able to infer landscapes across BPA concentrations in a single regression, assuming low-dimensional variation in community-function landscape coefficients. We developed a regularized optimization procedure to accomplish this and found that the variation in AUC across BPA concentrations enabled such an approach (Fig. S12).

### Functional Landscape Enables Design for BPA Remediation in Soils

All of the experiments described above were performed with synthetic communities in a minimal medium with BPA as the sole carbon source. In these conditions, organisms have replete nutrients (nitrogen, phosphorus, sulfur, etc.), and do not compete for resources with other strains. In a remediation context, organisms are often added to settings such as the gut [19, 20, 48] or soils [9, 49], where they compete with or displace an extant microbiome. A major problem with biological remediation is getting the inoculated microbes to engraft in a complex natural environment.

Recently, Silverstein *et al*. suggested that engraftment might be more successful when the inoculated community can strongly occupy a niche that is only weakly occupied, or unoccupied, by the native microbiome [23]. Motivated by this hypothesis, and the fact that BPA degradation is a metabolically complex process not readily performed by all bacterial taxa [50], we wanted to test whether our *in vitro*-designed BPA degrading consortia could drive BPA degradation in soils.

To accomplish this, we created BPA-contaminated soil slurries by mixing soil with BPA-contaminated water. We modified our BPA measurement method to enable extraction from these slurries (see Methods). We then inoculated each of the ten communities designed for a range of *in vitro* degradation performances (Fig. 3C) in slurries made with two different soil samples, each contaminated with two different BPA concentrations (100 and 125 ppm). As in the experiments above, we measured BPA concentrations in time via sampling (Fig. 6A). To assess the ability of the native soil microbiome to degrade BPA, we included a control where no community of isolates was added.

**Figure 6:**
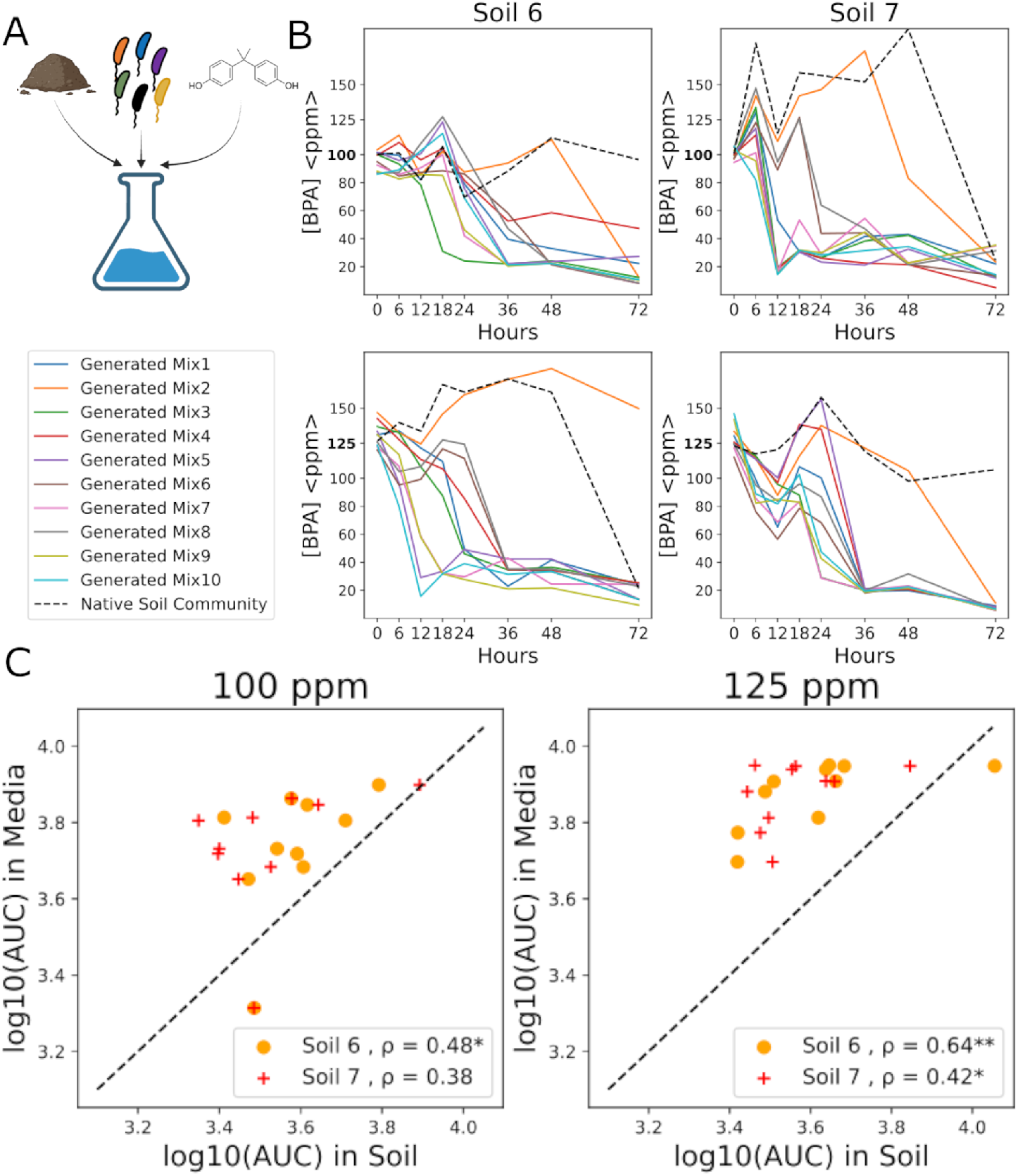
Designing for BPA Remediation in Soils. **(A)** Schematic for BPA remediation protocol. Soil was combined with BPA-contaminated water to form a slurry. Contamined slurries were inoculated with synthetic communities, and BPA degradation was assayed (see Methods). **(B)** BPA degradation curves for the ten synthetic communities in two soils and two initial BPA concentrations: 100 ppm (top) and 125 ppm (bottom). The black line shows BPA degradation by the native soil microbiome without an added synthetic community. **(C)** Scatter plots showing the ten communities’ AUC (Fig. 2) in soil (x-axis) and *in vitro* (y-axis) for two soils and two initial BPA concentrations. Dotted lines denote y=x. *ρ* in the legend denotes Pearson’s correlation between *in vitro* and in soil degradation of BPA. *p < 0.05, ** p < 0.01 determined from 100 bootstrapping iterations rejects the null hypothesis that the Pearson’s correlation coefficient is <0.

We found that adding communities to soils increased the rate of BPA degradation. In some cases, the native microbiome was not able to degrade BPA on its own (Soil 1, 100ppm, Soil 2, 125ppm). Furthermore, the BPA degradation dynamics in soils depended on which synthetic community was inoculated, suggesting that community composition is important for determining BPA degradation even in the presence of a complex indigenous microbiome. We conclude that designed synthetic communities can impact BPA remediation in soils.

Finally, given that our landscape models enable the rational design of degrading communities (Fig. 3) we wanted to know whether the performance of a given community *in vitro* provided any insight into the performance when the same community was used for remediation in soils. To accomplish this, we compared the AUC for each community *in vitro* with the AUC for BPA degradation by that community in soils. We note that given the challenges of extracting BPA from soil, we do not expect a 1:1 relationship between these two AUC values. However, we find that the AUCs *in vitro* correlate with the AUC in soils for both soils (Fig. 6). This result shows that the landscapes learned for synthetic communities in vitro are informative of community performance in soils. Our results establish an example of successful bioremediation by designing communities to strongly occupy a niche (in this case, BPA degradation) as posited by Silverstein *et al*. [23].

## Discussion

Using a library of BPA-degrading isolates, we showed that the community-function landscape of BPA degradation can be learned statistically from ensembles of communities where BPA degradation dynamics are quantified. Learning the landscape statistically required only knowledge of the strains present in the community, and we inferred these landscapes across a range of initial BPA concentrations. Inspection of these landscapes showed that as BPA concentration increased (and with it, toxicity), so did epistasis. We concluded that collective effects become more important as toxicity rises, and indeed monocultures cannot degrade BPA at the highest concentration. We then showed that what we learned from this landscape translated into the remediation of BPA in soil samples.

Functional landscapes are most easily inferred when they are relatively smooth with limited epistasis [15] or when global non-linear transformations render the landscape approximately additive [17]. Indeed, most of the previous work on landscapes falls into one of these two categories. The BPA degradation landscape presented here offers a slightly different picture: the landscape shows smooth, low-ruggedness at low BPA concentrations and higher ruggedness at high BPA levels (Fig. 4). However, we do observe a difference in model fits between the low and high concentrations, where higher concentrations were more difficult to fit (Fig. 3B). We attribute this difficulty to the sparse sampling of 70 communities (out of a possible *∼* 65, 000) in a regime where epistatic terms dominate the landscape (Fig. 5). Despite this difficulty, our success highlights an example of characterizing a rugged community function landscape.

Previous explorations of community-function landscapes are mostly limited to a stable environment and exploring changes in community composition [15, 19, 51]. Our results demonstrate the power of functional landscapes as a tool for understanding environmental changes. In particular, we showed that the functional landscape changes with initial BPA concentration smoothly, to the extent that `SoftImpute` accurately predicts BPA degradation of communities (Fig. 2D) and that regression coefficients vary in a low-dimensional manner (Fig. S7). Our work serves as a proof of principle for analyzing community function landscapes across other environmental gradients or even multiple environmental gradients simultaneously (such as the effect of pH on BPA degradation landscape). The low-rank regressor formalism developed here (Fig. S12) serves as a generic method for inferring landscapes across environmental gradients.

We showed that the smooth variation in the landscape is accompanied by epistasis that increases with BPA toxicity. At low BPA concentrations, the presence of degraders is the primary driver of degradation, with statistical interactions between them generally competitive (Fig. 5). The importance of epistatic interactions for BPA degradation rises with toxicity, where at the highest BPA concentration measured, no individual strain can degrade BPA (Fig. 4 D). The importance of epistatic interactions in our landscapes is especially evident in the community of all strains - Mix20 - not being the fastest to degrade BPA in all five concentrations measured (Fig. 1C). This result shows that composition matters in a non-trivial manner. The observation is consistent with the presence of many instances of both statistical cooperation (negative *γ* coefficients, speeding up degradation) and competitive (positive *γ* coefficients, slowing down degradation) effective interactions between strains. Our work provides unique insight into the power of a functional landscape in determining epistatic interactions that are biologically relevant.

Previous efforts in bioremediation have encountered difficulties in engrafting bacteria into soils [52, 53], resulting in low bioremediation efficiency [24]. Some studies have attempted to model complex interactions between invaders and resident consortia to design communities for remediation [7]. Our work highlights a practical application of the theory posited by Silverstein *et al*. [23] in combination with community function landscapes to address this problem. This success paves the way for simplifying similar endeavors for other bioremediation targets.

## Methods

### BPA Media

BPA medium was prepared using 1x M9 medium with BPA as the sole carbon source (Tbl. S1). BPA was dissolved in 1x M9 medium and autoclaved for 30 minutes at 121°C. The final concentration of the medium was verified by the assay described below before use.

### Soil Sampling and Enrichment

Five sites were identified as being potentially BPA contaminated—two directly adjacent to BPA-producing factories, and three directly downstream from these factories (Tbl. S2, sites 1 —5). Approximately 2 kg of topsoil from each site was harvested and refrigerated at 4 °C for approximately one month before use. Soil enrichments were performed in 30 ppm BPA M9 media. Enrichments were performed in 250 ml Erlenmeyer flasks, containing 3 g of soil in 60 mL of BPA media. 400 µg/mL of cycloheximide and 40 µg/mL of nystatin were added to inhibit eukaryotic growth. The solution was vortexed, and a 40 µL sample was taken for initial BPA concentration measurement. Flasks were incubated aerobically at 30°C, shaking at 220 RPM.

BPA concentration was measured daily. Whenever the measured BPA concentration fell below half of the starting concentration, the sample would be diluted into fresh BPA media adjusted for a final volume of 60 ml, up to three rounds of dilution. The dilution factor increased with each round —the first was 1:16 (3.75 mL sample with 56.25 mL BPA media), then 1:32, then 1:64. The purpose of dilutions was to select for microbes that either degrade BPA or co-exist with degraders in the presence of BPA.

From soils that degraded BPA after the third dilution, samples were diluted in PBS to a 10^*-*5^ dilution and plated on R2A plates (R2 Broth, Research Prodcuts International, product number R24260; Agarose, Fischer Scientific, product number BP1356). Plates were incubated at 30 °C to encourage growth. Colonies were picked and then streaked to purity, after which isolates were mixed 1:1 with 50% glycerol and stored at -80°C. Isolates were sequenced via Sanger 16s sequencing using the universal 16S primers 27F (AGAGTTTGATCMTGGCTCAG) and 1492R (TACGGTTACCTTGTTACGACTT)

### Individual Strains

16 strains were isolated and identified from soil enrichments. Ten of these strains degraded 30 ppm BPA on their own, while six did not. Our BPA degrading strains came from two soil samples, while our nondegraders came from the other three samples. 16S Sanger sequencing was performed for each strain (as mentioned above), and DNAStar Lasergene 17 [54] was used to construct consensus sequences. Strain taxonomy was identified through NCBI BLAST [55]. The EMBL-EBI Job Dispatcher [56] was used to align sequences via Muscle and construct the phylogenetic tree (Fig. ?? B) and the Interactive Tree of Life [57] was used to visualize and edit the tree.

### Measuring CO_2_

CO_2_ production in our ten degrading isolates by BPA Degradation was measured by a Microresp assay (Fig. S3). A monoculture of each strain was prepared as described in Methods (‘Synthetic Community Experiments’). Four technical replicates were prepared: two in 60 ppm BPA media, and two in M9 media with no carbon source. 700 µL of each replicate was transferred into a 96-deep well plate, which was then sealed with a pH-sensitive dye in an agarose gel. Respiration of CO_2_ reduces the pH of the dye, thus changing its color (See de Jesús Astacio *et al* [58] for more information on the setup of the Microresp system). The plate was incubated at 30°C shaking at 900 RPM for three days, after which CO_2_ respiration was measured by an absorbance reading of the dyes.

### BPA Assay

BPA concentrations were measured using a modified version of the phenol colorimetric assay [59] optimized for high-throughput use in a 96-well plate. Under basic pH conditions, 4-Aminoantipyrine (4-AAP) reacts with BPA in an electrophilic substitution reaction, which can then be oxidized by potassium ferricyanide to produce a red dye that absorbs at 506 nm in proportion to the initial BPA concentration in the sample. Reagents were added in the following order: 120 µL of 0.25M sodium bicarbonate (Sigma-Aldrich, product number S6014) to serve as a basic buffer, 40 µL of a BPA containing sample sample, 40 µL of 20.8 mM 4-AAP (Sigma-Aldrich, product number A4382), and 40 µL of 83.4 mM potassium ferricyanide (Arcos Organics, product number 424125000). The plate is incubated at room temperature, shaking at 750 RPM for 45 seconds, and then incubated statically for ten minutes. Afterward, the absorbance of each sample was read at 506 nm. A standard curve was made from solutions of known BPA concentration in BPA minimal media as described above for every timepoint in every experiment, and each 96-well plate was prepared with its own standard curve. The standard curves had a slope of 133.36 ±13.14 and an intercept of -0.88 ±3.1(Fig. S1).

### Synthetic Community Experiments

To prepare our strains for mixing into synthetic communities, each of the strains was sampled from its frozen stock and grown anaerobically in 5 ml Tryptic Soy Broth (TSB) media for 44 hours in tightly-closed falcon tubes, incubating at 30°C and 220 RPM. After incubation, samples were centrifuged at 4000 RPM for 15 minutes and the supernatant was pipetted out. Pellets from each sample were resuspended in 1ml M9 media as a wash step, and centrifuged again at 4000 RPM for 8 minutes, after which the supernatant was pipetted out again. Finally, pellets were suspended in 2 ml of BPA media, and samples were ready for community assembly.

All communities were mixed and incubated in Fischer 48 deep well plates (part number 1162B95), with one well housing one community. Each well was filled with 2 mL of M9-BPA medium, and each isolate was added at an initial nominal OD of 0.02 irrespective of the community richness. All wells were mixed thoroughly by pipetting up and down repeatedly and then incubated at 30°C in a Fisher Scientific benchtop microplate shaker (Catalog number 02-217-757) shaking at 900 RPM, covered by a breathe-easy membrane (Sigma-Aldrich, part number Z380059). Every plate also contained at least one well with no bacteria added to it (as an evaporation control) and one well with TSB medium (as a contamination/splashing control). Every community was run in technical duplicates.

Samples were taken at time points 0, 6, 12, 18, 24, 36, 48, and 72 hr, and the BPA concentration was measured for each sample at each time point as previously described. Positive controls ensured degradation dynamics across different batches of experiments were comparable, while negative controls detected contamination and also served as evaporation controls.

### Imputation to predict AUCs across concentrations of BPA

In Fig. ?? we predicted community AUCs via matrix imputation. Imputing entries from our AUC matrix Z (Fig. ??) was done via the R package `SoftImpute`. Every entry of *Z* is denoted *z*_*c,k*_, which is the *log*_10_(*AUC*) of community c in concentration *k*. We iteratively dropped out every entry of the matrix (along with all technical replicates), to achieve *Z*_*-c,k*_ with community *c* in concentration *k* dropped out. We then process *Z*_*-c,k*_ to be 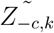, where every column *k* is *z*-scored. and fit `SoftImpute` using the 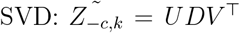, where *D* is soft-thresholded (*D*^*^)to minimize its nuclear norm. The *Z* matrix is reconstructed by D^*^ as *Z*^*^ = *UD*^*^*V* ^⊤^, where 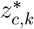 is the imputed value of the dropped entry. Leave-one-out cross-validation was done to choose the soft-thresholding regularization parameter over a range of 40 values between 0 and 10 which maximizes the total R^2^ of predictions (Fig. S6 A). For all fits, the resulting D^*^ was rank 2.

We also performed a similar imputation by randomly dropping out 10% of the entire matrix (accounting for technical replicates), repeated over 100 iterations of 10% dropout (Fig. S6 B).

### Linear Regressions

Regressions were performed using the Python `scikit-learn` package. Ridge was chosen over LASSO because the *L*_2_ penalty decreases all coefficient values to avoid overfitting, while *L*_1_ regularization performs variable selection which can be problematic when regressors are correlated [60]. For each concentration *k*, models were fit of the form

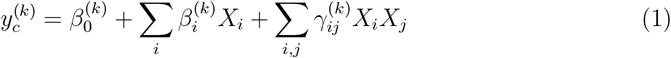

where 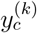 is the *z*-scored *log*_10_(*AUC*) of community *c* in concentration *k* (similar to our imputation method above, 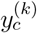 is *z*-scored within each model for concentration *k*), 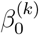 is the intercept that is fit from the data for concentration *k, X*_*i*_ represents the presence and absence of strain *i* in the community *c*, denoted in +1/-1 notation to simplify interpreting coefficients [15, 18, 61], 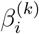 is the additive coefficient for stain *i* in the model fit from concentration *k*, and 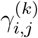 is the pairwise coefficient for strains *i* and *j* together in the community *c* in the model fit from concentration *k*. The objective function for each model was

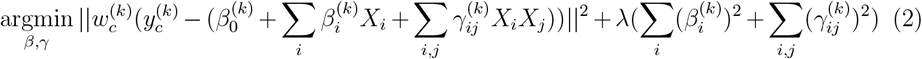

where 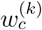 is the weight applied to community *c* in the model for concentration *k* and λ is the *L*_2_ penalty term. Weights were determined through a Gaussian Kernel Density Estimate of community AUCs using `scikit-learn`. For each 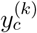, the estimator computes

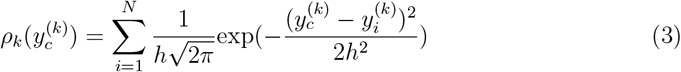

where 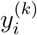 represents all measured *y*^(*k*)^, and *h* represents the bandwidth of the curves. For each concentration, a grid search of *h* between 10^*-*3^ and 1 was performed to find the bandwidth that maximized the log-likelihood of the estimated distribution of all 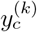. For all other parameters, the default `scikit-learn` settings were used. The final weight 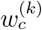 was set to be 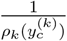 so that a community whose AUC is more ‘rare’ within concentration *k* are given greater weight in the regression.

**Table 1:** Ridge Models Optimal Penalty Terms

| Concentration | Value for L2 Penalty Term |
| --- | --- |
| 60 ppm | 82.07 |
| 80 ppm | $1.10 \times 10^{-5}$ |
| 100 ppm | 133.57 |
| 125 ppm | 32.41 |
| 150 ppm | 16.64 |

The optimal λ was chosen through leave-one-out cross-validation over a range of 10,000 values between 10^*-*5^ and 10^10^ and finding the value that minimized the objective function above (similar to our imputation method above, z-scoring of 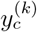 was only performed after excluding the validation community). Once the optimal λ was identified, the predicted value for each community within each concentration was plotted against the measured value (Fig. 3 B). A separate model was fit to all *y*^(*k*)^ within each concentration *k*, each with their own optimal λ (Tbl. 1). The cross-validation for the model fit to data from 80 ppm BPA yielded the smallest tested value for λ (near 0), in stark contrast to the other four models. The strong model fit (Fig. 3 B) indicates the general ease of fitting the model to this data compared to data from the other four initial concentrations.

### PCA Analysis

PCA analysis was performed on our matrix of regression coefficients (Fig. S7 C) to determine the variation in community function landscapes. For each concentration *k*, the column of our matrix becomes 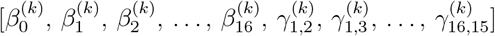 The coefficients were scaled by subtracting the mean and normalizing by the standard deviation within each concentration before being subject to PCA transformation.

### BPA Extraction and Community Inoculation in Soils

Two soils were sampled in amounts of roughly 2 kg from prairie fields (Tbl. S2, sites 6 & 7). Soils were sieved through to 2mm, and a portion of each was sterilized by autoclave at 121°C for 99 minutes three times, with 24 hours in between each time and the last occurrence being 8 hours before preparing the communities. 0.85 g of each soil was suspended in 1.7 mL of 100 ppm BPA media in 48 deep well plates and vortexed to ensure sufficient mixing of BPA. Strains were added in an identical manner to community mixing experiments. Incubation was also performed similarly. For each soil, a sample was prepared with no community added, and another sample with non-BPA media for controls. Two technical replicates for each condition were prepared. To measure BPA concentration at each sampling time point, BPA needed to be extracted from the soil slurry. 100 µL of the slurry was sampled and mixed with 150 µL of 100% ethanol. The samples were then vortexed and then spun down at 4000 RPM for 2 minutes. 40 µL of the supernatant was sampled to use for our BPA concentration assay, with an extraction efficiency of approximately 60% (Fig. S10).

## Supporting information

Supplemental Figures and Analysis

## Acknowledgments

We thank members of the Kuehn laboratory for guidance and discussion and members of the Raman laboratory for assistance with the protocol for synthetic community experiments. We thank Dr. Vaibhhav Sinha for his assistance with the Microresp CO_2_ protocol. M.Y. acknowledges support from the Department of Education under grant number P200A210054 and the National Institute of Health under grant number T32GM007281. S.K. acknowledges the National Institute of General Medical Sciences R01GM151538, and support from the National Science Foundation through the Center for Living Systems (grant no. 2317138). S.K. and R.W. acknowledge financial support from the National Institute for Mathematics and Theory in Biology (Simons Foundation award MP-TMPS-00005320 and National Science Foundation award DMS-2235451). Any opinions, findings, conclusions, or recommendations expressed in this material are those of the authors and do not necessarily reflect the views of the National Science Foundation.

## Author contributions

M.Y., K.K.L. and S.K. conceptualized the research. M.Y., K.K.L. and S.K. designed the experiments. M.Y. and J.T. conducted field sampling, enrichments experiments, and isolations. M.Y. performed BPA assay optimization, community assembly experiments, and the all statistical analysis of the data except the low-rank regression (LRR) analysis. V.C. and R.W. designed the LRR algorithm and provided the code. S.K. supervised all aspects of the study. M.Y. and S.K. wrote the manuscript.

## Data Availability

All datasets generated in our study are available within the following Open Science Framework (OSF) project: https://osf.io/gcfr8/. Some datasets are available within the Supplementary Tables.

## Code Availability

All code written for models and analysis was written in either python or R, and is available at https://osf.io/gcfr8/

## Notes

https://osf.io/gcfr8/

### Competing Interest Statement

The authors have declared no competing interest.

