## Supplemental Figures and Analysis for "Collective Microbial Effects Drive Toxin Bioremediation and Enable Rational Design"

Table S1: M9 Media Components

| Reagent | Amount |
|---|---|
| 5X M9 Salts (BD Biosciences, product number 248510) | 5.64 g |
| 1M MgSO4 (Fischer Scientific, product number AA3333736) | 1ml |
| 1M CaCl2 (Fischer Scientific, product number 0556) | 50 ul |
| MiliQ Water | Remaining volume up to 500 ml |
| BPA (Sigma-Aldrich, product number 133072) | As needed |

Table S2: Table for Soil Sampling Sites

Table S3: Data Regarding 16 Strains

### Supplementary Information

#### Regressions on Random Shuffles of log10(AUC)s

To evaluate the significance of our model fits (Fig. 3) and all resulting analyses, we sought to employ a null model of random AUCs for each community. To do this, within each initial BPA concentration $k$, we de-coupled $y_c^k$ from community composition $c$ by randomly shuffling the order of the response variable $y_c^k$, This randomization enforces no relationship between a community composition and associated AUC. After random shuffling, models were fit in the same way as described in Methods. The weighting scheme was not changed; therefore, every $y_c^k$ retained its weighting $w_c^k$.

For each model, 100 random shuffles and subsequent fits were performed, and $\rho$ (Pearson's) between predicted and "true" AUC were compared to that of the original models. The data (Fig. S5B) shows that models fit to random shuffles of the response variable perform significantly worse than the original models, where $p$-value $< 0.01$ for the lower three concentrations and $p$-value $< 0.05$ for the higher two concentrations. The result suggests that our linear models are utilizing community composition to predict AUC in a statistically significant fashion.

#### Increasing model order

Skwara *et al.*[15] observed that for low-ruggedness landscapes, first-order models explained most of the variance in the functions measured, while second-order coefficients improved model fits and third-order coefficients mostly did not improve model performance. However, the landscapes measured here are more rugged than those encountered by Skwara *et al.* therefore we wanted to test whether the same observations hold for this landscape.

We fit AUC data from all five initial BPA concentrations into two additional models of the form Eqn. 1: one only including first-order terms, and one including third-order coefficients alongside first- and second-order terms. The model including only first-order terms was:

$$y_c^{(k)} = \beta_0^{(k)} + \sum_i \beta_i^{(k)} X_i \tag{4}$$

while the model including third-order terms was:

$$y_c^{(k)} = \beta_0^{(k)} + \sum_i \beta_i^{(k)} X_i + \sum_{i,j} \gamma_{ij}^{(k)} X_i X_j + \sum_{i,j,k} \zeta_{ijk} X_i X_j X_k \tag{5}$$

All models were fit in the same way and regularized with the $L_2$ penalty (see Methods), and model fits were compared by comparing the resulting RMSEs of leave-one-out cross-validation.

Results (Fig. S5 C) show us that, on average, adding third-order terms (green) does not substantially improve model fits compared to a model up to second-order coefficients (orange), with the only major difference seen at 125 ppm. However, we do see that fitting the data to only first-order coefficients (blue) does not lose significant predictive power compared to the second-order model, with the models fit to 100 ppm and 150 ppm data being nearly equivalent. The only major difference is seen at 60 ppm, the least epistatic model of the five, in which second-order coefficients greatly improve model performance.

Naively, it may appear that the results of Fig. S5C contradict the notion that epistasis increases with increasing BPA concentration. However, we note that the quality of the fit (RMSE, y-axis Fig. S5C) does not indicate the *relative importance or magnitude* of additive vs epistatic coefficients in the regression [62]. This is quantified via the F1-statistic (main text).

#### $F_1$ statistic In-Silico Experiments

The $F_1$ statistic (Fig. 5) indicates that overall epistasis increases as BPA toxicity increases. However, we also note that the regression fits worsen as the initial BPA concentration increased (Fig. 3B). Although BPA degradation data from monocultures and communities confirms the increased importance of a community to degrade BPA at higher concentrations, we wanted to ensure that the decrease in the $F_1$ statistic was not an artifact of the worsening model fits with BPA concentration.

To systematically investigate this, we considered a system of $N = 10$ binary variables representing strains, denoted as $x_i \in \{-1, 1\}$ for $i \in \{1, \ldots, N\}$. $x_i^c$ defines the presence or absence of strain $i$ in community $c$. For each community $c$, we defined the corresponding functional output $y_c$ using a second-order model of the same form as Eqn. 1:

$$y_c = \sum_{i=1}^{N} \beta_i x_i^{(c)} + \sum_{i=1}^{N} \sum_{j>i}^{N} \gamma_{i,j} x_i^{(c)} x_j^{(c)} + \eta_c \tag{6}$$

where $y_c$ is the functional output of community $c$, $\beta_i$ and $\gamma_{i,j}$ are additive and pairwise coefficients respectively (see Methods), and $\eta_c \sim \mathcal{N}(0, \sigma^2)$ is independent Gaussian noise with variance $\sigma^2$.

We generated a family of synthetic models and controlled the true $F_1$ statistic of the model to be between 0.1 and 1, in increments of 0.1. This was achieved by varying the proportion of additive ($\beta_i$) versus epistatic ($\gamma_{i,j}$) coefficients.

We first sought to evaluate how the number of sampled communities affects the estimated $F_1$ statistic. For each model $f$ (corresponding to a fixed $F_1$ statistic), we generated $S = 60$ communities, denoted as $X^S$. We then sampled from $X^s$ a number of communities $s'$, varying from $s' = 5$ to $s' = 55$. For each sampling size $s'$, we generated $R = 100$ independent random subsamplings. Each subsampling was denoted $X_r^{s'}|_{r=1}^{R}$. We also added a sample for all 60 communities, i.e. $s' = 60$. We then computed functional outputs for the sampled communities, $y_c^{s',r} = f(x_c^{s',r})$ for each community $c^{s',r} \in X_r^{s'}$. (for now, $\sigma^2 = 0$ in $f$, thus $\eta_c = 0$ for all communities $c$.). From the complete dataset $\{X_r^{s'}, y_c^{s',r}\}$, we fit a regression model in the same way as the main analysis (see Methods) and inferred coefficients $\hat{\beta}_i$ and $\hat{\gamma}_{i,j}$. We used these inferred coefficients to calculate the inferred $F_1$ statistic. This was repeated for all sample sizes $s'$ and for all 100 random subsamplings of $X^{s'}$, such that for each $F_1$ value of $f$, a distribution was generated for each inferred $F_1$ statistic for each sample size $s'$, denoted $\hat{F}_1^{s',r}$. The mean of all 100 subsamplings was reported as a final $\hat{F}_1^{s'}$.

By systematically varying the number of sampled communities $s'$, we assessed how well $\hat{F}_1^{s'}$ approximates the true landscape ruggedness $F_1$. This allowed us to quantify the effect of sample size on the accuracy of the estimated ruggedness metric and determine whether certain landscape properties (e.g., degree of epistasis) influence this relationship.

Figure S11 A shows the results of this test. As expected, we find that when all 60 communities are used in the fitting, $\hat{F}_1^{s'=60}$ closely matches that of the underlying model. However, as the sample size $s'$ decreases, $\hat{F}_1^{s'}$ quickly becomes inaccurate. This effect is most dramatic in landscapes that are mostly or purely additive (i.e. $F_1 \approx 1$). Therefore, we conclude that highly rugged landscapes are not very susceptible to the effects of undersampling on the $F_1$ statistic of the inferred model.

We then wanted to ensure that the decreased $F_1$ statistics of the models fit to data from 125 and 150 ppm BPA are not explained by the poor model fits. We generated a single dataset $X^s$ of size 60 communities, and then generated a single regression function $F$ (of the same functional form as Eqn. 6 with fixed $\beta_i$, $\gamma_{i,j}$) which had an $F_1$ statistic of 0.168. This time, we varied $\sigma^2$, the variance of the noise in $\eta_c$, to be between $\sigma'^2 = 0.0625$ and $\sigma'^2 = 5$. We generated $R = 100$ instances of noise generated from $\eta_c$ for each community

$c$, and then calculated $y_c^{\sigma'^2,r} = F(x_{c^s})|_{r=1}^{R}$. Since the response variables were of the order —1, higher values of $\sigma'^2$ represented progressively worse model fits.

Regression models were then fit to all 60 communities in $X^s$ and all $y_c^{\sigma'^2,r}$, and the inferred $\hat{F}_1$ statistic was compared to that of $F$ (0.168).

The results (Fig. S11 B) show that on average, the $F_1$ statistic *increases* as a result of poor model fits, rather than decreases. This indicates strongly that the decreased $F_1$ statistic seen in the 125 and 150 ppm models (Fig. 5) are not due to the difficulty in fitting the underlying landscape.

#### Low Rank Regressor

Inspired by the success of `SoftImpute` as well as the low-dimensional structure of community AUCs (Fig. 2 D), we sought to pursue a regression model that would exploit the low-dimensionality of our data to inform the regressions.

The low-rank regressor is a model that attempts to learn coefficients from all initial BPA concentrations simultaneously. It assumes the relationship

$$Z \approx X\Theta Y, \quad \text{for an approximately low-rank matrix } \Theta \in \mathbb{R}^{p\times r}, \tag{7}$$

where $Z \in \mathbb{R}^{n_{\text{comm}}\times n_{\text{BPA}}}$ is the matrix of community AUCs under all available $n_{\text{BPA}}$ concentrations, $X \in \{\pm 1\}^{n_{\text{comm}}\times p}$ is the community composition matrix and $Y \in \mathbb{R}^{r\times n_{\text{BPA}}}$ is a polynomial basis matrix, with each row comprising monomials evaluated at the available BPA concentrations:

$$Y = \begin{bmatrix} 1 & 1 & \dots & 1 \\ \phi_1(\mathsf{PPM}_1) & \phi_1(\mathsf{PPM}_2) & \dots & \phi_1(\mathsf{PPM}_{n_{\text{BPA}}}) \\ \phi_2(\mathsf{PPM}_1) & \phi_2(\mathsf{PPM}_2) & \dots & \phi_2(\mathsf{PPM}_{n_{\text{BPA}}}) \\ \vdots & & & \vdots \\ \phi_r(\mathsf{PPM}_1) & \phi_r(\mathsf{PPM}_2) & \dots & \phi_r(\mathsf{PPM}_{n_{\text{BPA}}}) \end{bmatrix}. \tag{8}$$

Here, $r$ is the degree of the polynomial, $\phi_1, \dots, \phi_r$ are the elements in the polynomial basis, and $\mathsf{PPM}_j$ denotes the $j^{\text{th}}$ BPA concentration. In our experiments, we choose $\{\phi_j\}$ to be the monomials, with $\phi_k(x) := x^k$; however, alternative choices may be able to provide better fits. Under this formalism, the product $\Theta Y \in \mathbb{R}^{p \times n_{\text{BPA}}}$ is the matrix of coefficients for all observations and satisfies $\text{rank}(\Theta Y) \leq r$ by construction.

**Fitting**

We calculate an estimate $\widehat{\Theta}$ from data by solving the following optimization problem:

$$\widehat{\Theta} := \underset{\Theta}{\text{argmin}} \, \|W \odot (Z - X\Theta Y)\|_{\text{F}}^2 + \lambda_1 \|\Theta\|_* + \lambda_2 \|\Theta\|_{\text{F}}^2 , \tag{9}$$

where $W \in \mathbb{R}_+^{n_{\text{comm}} \times n_{\text{BPA}}}$ is a nonnegative weighting matrix (see Methods for our weighting scheme) and:

- $\odot$ denotes the elementwise (Hadamard) product;
- $\|\Theta\|_* := \sum_{i=1}^{r} \sigma_i(\Theta)$ is the *nuclear* norm, equal to the sum of all singular values of $\Theta$;
- $\|\Theta\|_{\text{F}} = \sqrt{\sum_{i,j} \Theta_{i,j}^2}$ is the so-called *Frobenius* norm;
- $\lambda_1$ and $\lambda_2$ are nonnegative, tunable parameters controlling the regularization strength.

The sum of the last two terms in (9) is a regularizer akin to the well-known Elastic Net [63]. The nuclear norm acts as an $\ell_1$ penalty on the singular values, promoting low rank in the solution $\widehat{\Theta}$, while the squared norm induces shrinkage in the learned coefficients like an $\ell_2$ penalty.

The optimization problem (9) is convex (in fact, *strongly* convex owing to the presence of the squared Frobenius norm). As such, it is amenable to several convex optimization solvers. For our purposes, we use the `SCS` solver [64] via its Python interface available in `cvxpy` [65]. The `SCS` solver is a general-purpose conic optimization solver: it first transforms (9) into an equivalent conic representation (which typically yields a constrained optimization problem) and applies an operator splitting method with algorithmic embellishments, including a variant of Anderson Acceleration [66], to solve the resulting optimization problem. The `SCS` interface provides access to several options that control its behavior during a solve; our code uses the default settings[1] except for the relative and absolute feasibility tolerances, $\epsilon_{\mathsf{rel}}$ and $\epsilon_{\mathsf{abs}}$: we set these to $\epsilon_{\mathsf{rel}} = \epsilon_{\mathsf{abs}} = 0.005$.

##### Model selection

We use cross-validation across BPA concentrations to select the regularization coefficients $\lambda_1$ and $\lambda_2$. Each regularization coefficient was scanned across 20 values logarithmically spaced between 0.001 and 1000. In particular, for each candidate $(\lambda_1, \lambda_2)$ pair, we iteratively select one row of $Z$ to act as the "held-out" community, use the remaining rows to fit $\widehat{\Theta}$ by solving (9) with the given $(\lambda_1, \lambda_2)$ pair, and compute the RMSE score of the fit on the held-out community; finally, we compute the average RMSE scores over held-out communities (Fig. S12 bottom right panel). We select the $(\lambda_1, \lambda_2)$ pair that achieved the lowest average score and retrain the model using all available communities. The optimal regularization coefficients were $(\lambda_1 = 4.833, \lambda_2 = 784.760)$, and the resulting $\widehat{\Theta}$ was rank-2.

Fig. S12 shows the fit of the LRR to the five BPA concentrations. In comparison to our standard linear regressions (Fig. 3), within each individual concentration, the LRR slightly underperforms the linear regression. However, the LRR has two constraints not shared with the linear regression: 1) the LRR only fits complete matrices $Z$ and $X$ and therefore leave-one-out cross-validation can only be performed by dropping a community out of all five concentrations simultaneously versus one-by-one, 2) the low-rank nature of $\widehat{\Theta}$ restricts the potential coefficients for all five concentrations, whereas the linear

[1] See `https://www.cvxgrp.org/scs/api/settings.html` for a list of available settings.

regression is free to have different parameters for each of the five concentrations. Given these restrictions, the fit of the LRR to the data is surprisingly well.

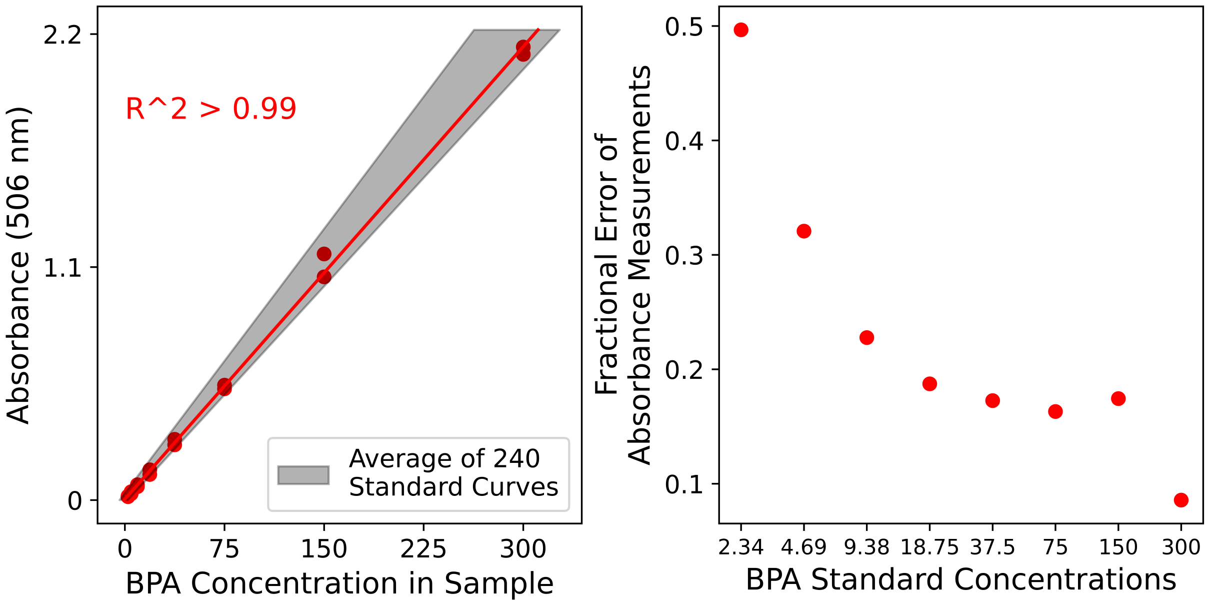


Figure S1: **Standard Curve of BPA Concentration Assay (LEFT)** A sample standard curve of BPA concentration, along with the line of best fit (red). $R^2 = 0.9998$ as calculated by the `scikit-learn` function `r2_score`. Gray shaded region indicates the error of the slope of the average standard curve (see Methods). **(RIGHT)** The fractional error of absorbance measurement as a function of the prepared BPA standard concentration. Fractional error is large at the lowest concentrations and small at the highest concentrations.

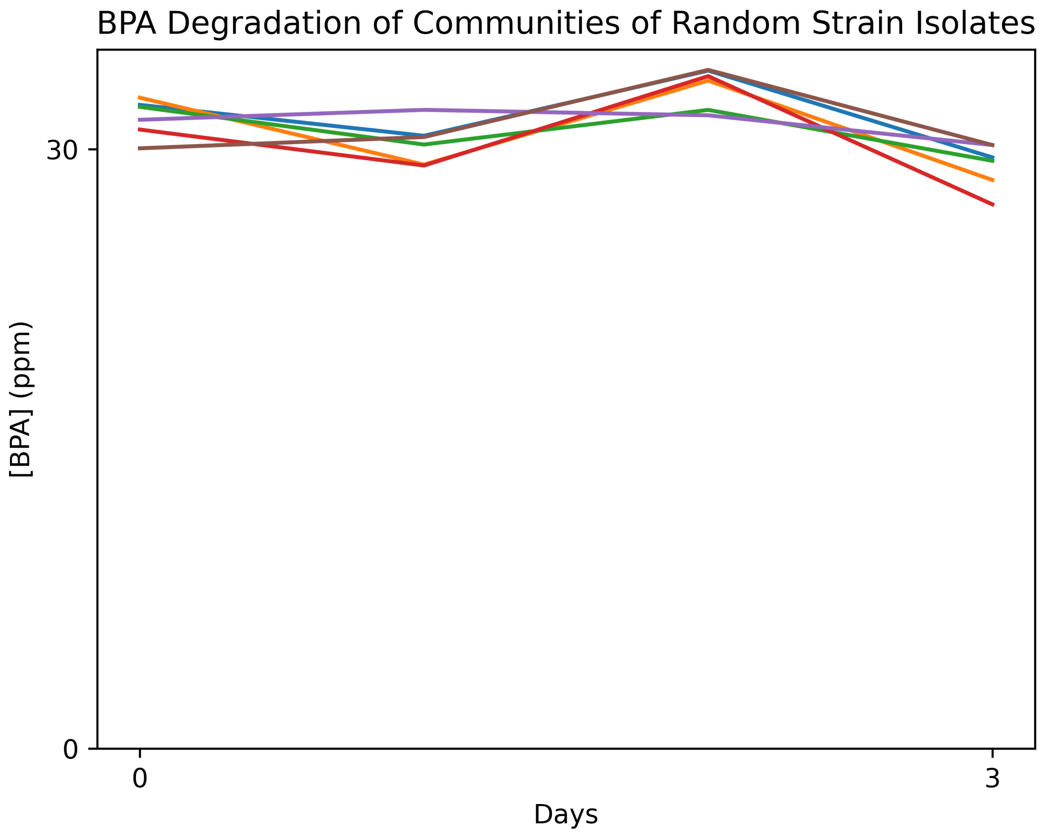


Figure S2: **Random Communities from Soil Isolates Do Not Degrade BPA** 5 random communities were assembled from six Proteobacterial strains isolated from various forests and plains within Illinois. These strains have been previously characterized for denitrification [44]. Communities were inoculated in 30 ppm BPA media, and the BPA concentration was measured every 24 hours for each community as described in the Methods section.

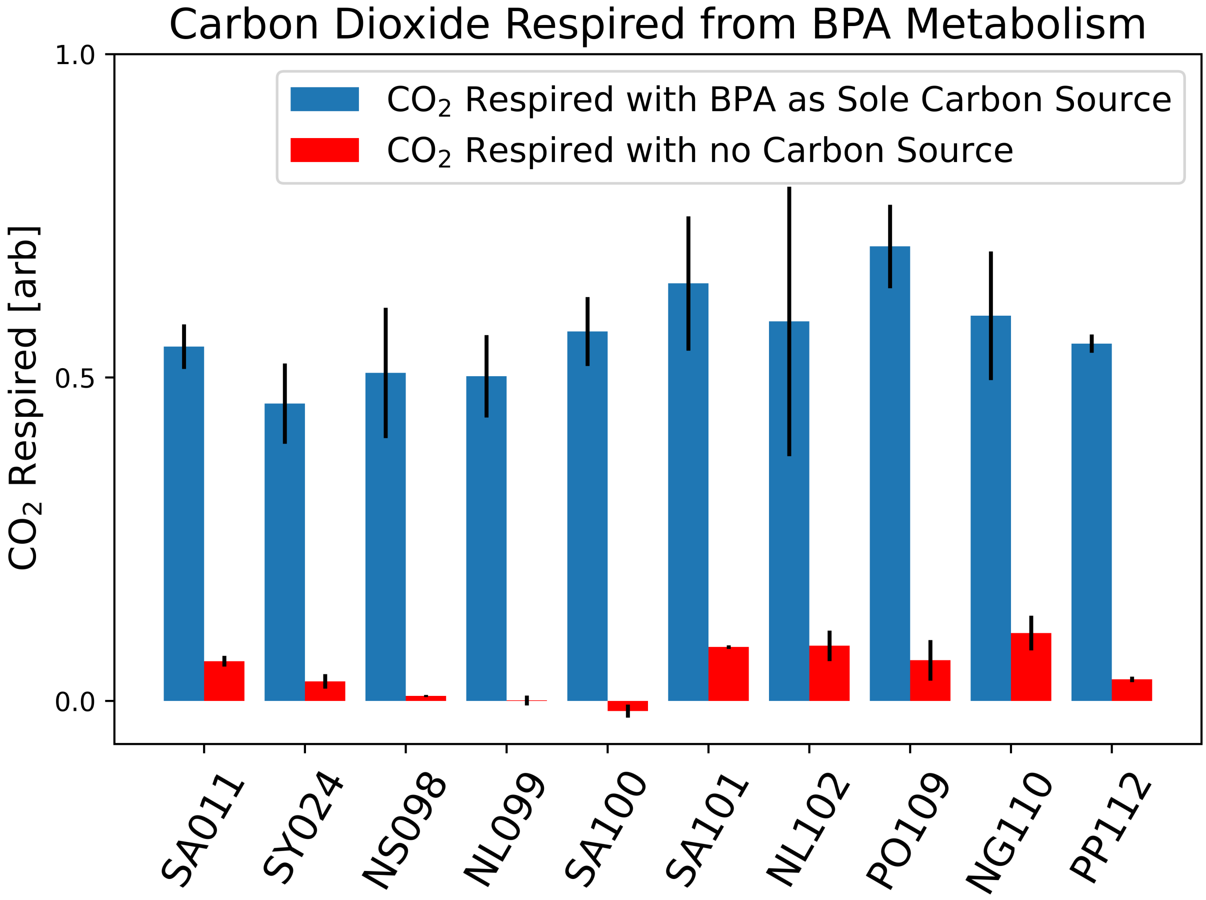


Figure S3: **$CO_2$ production by three strains during BPA degradation** Net $CO_2$ production by the 10 BPA-degrading strains when individually inoculated in 60 ppm BPA (blue) and when inoculated in minimal media with no carbon source (red) for three days. Error bars represent the standard deviation of technical duplicates. $CO_2$ measurements were performed with the microresp platform as described in the Methods section.

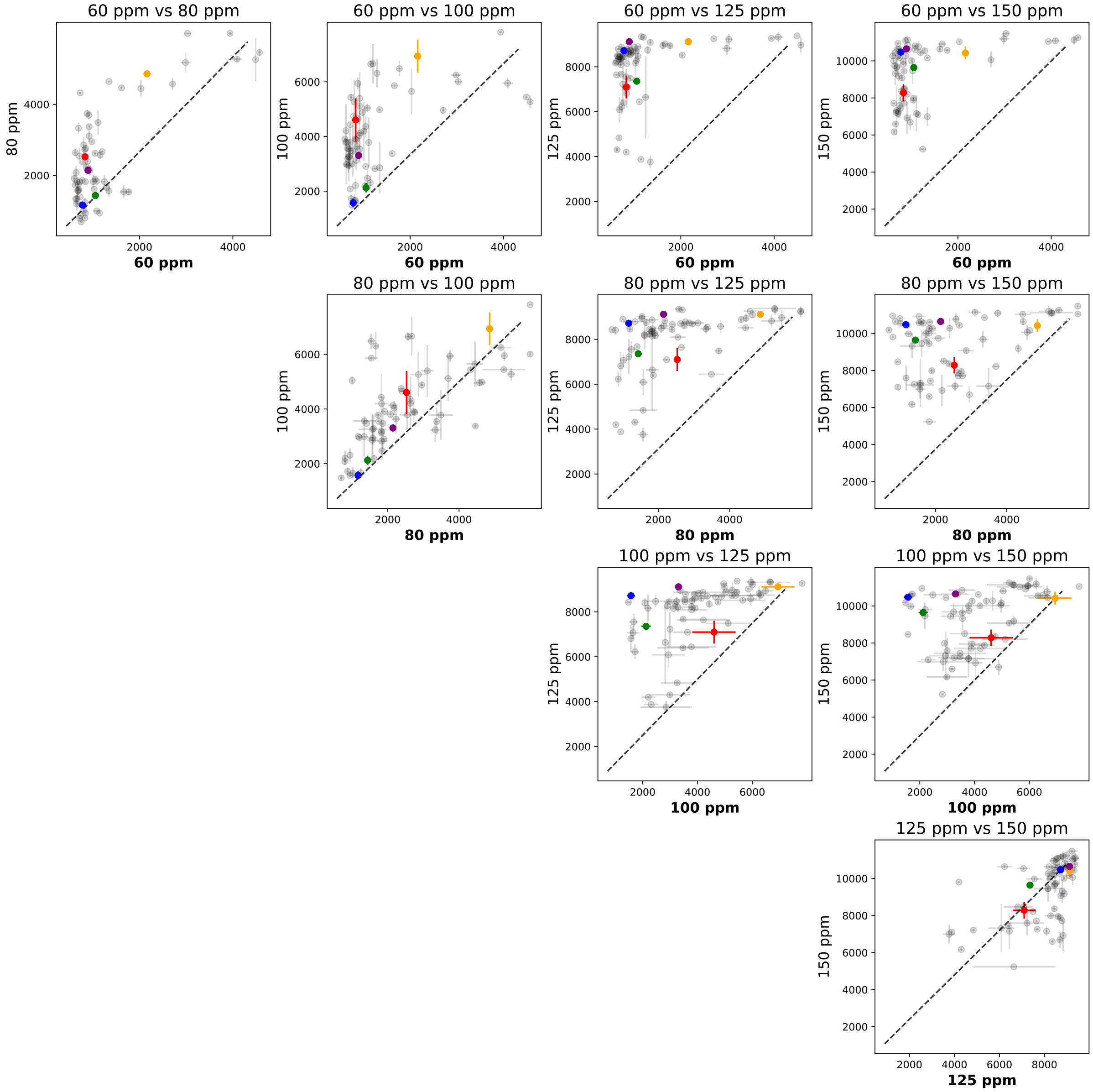


Figure S4: **Comparing Degradation AUC Across the 5 Initial Concentrations** Comparing the AUCs of our 70 communities across the five different initial concentrations. Error bars represent the standard deviation of technical replicates. Dotted lines indicate normalized equivalent AUC.

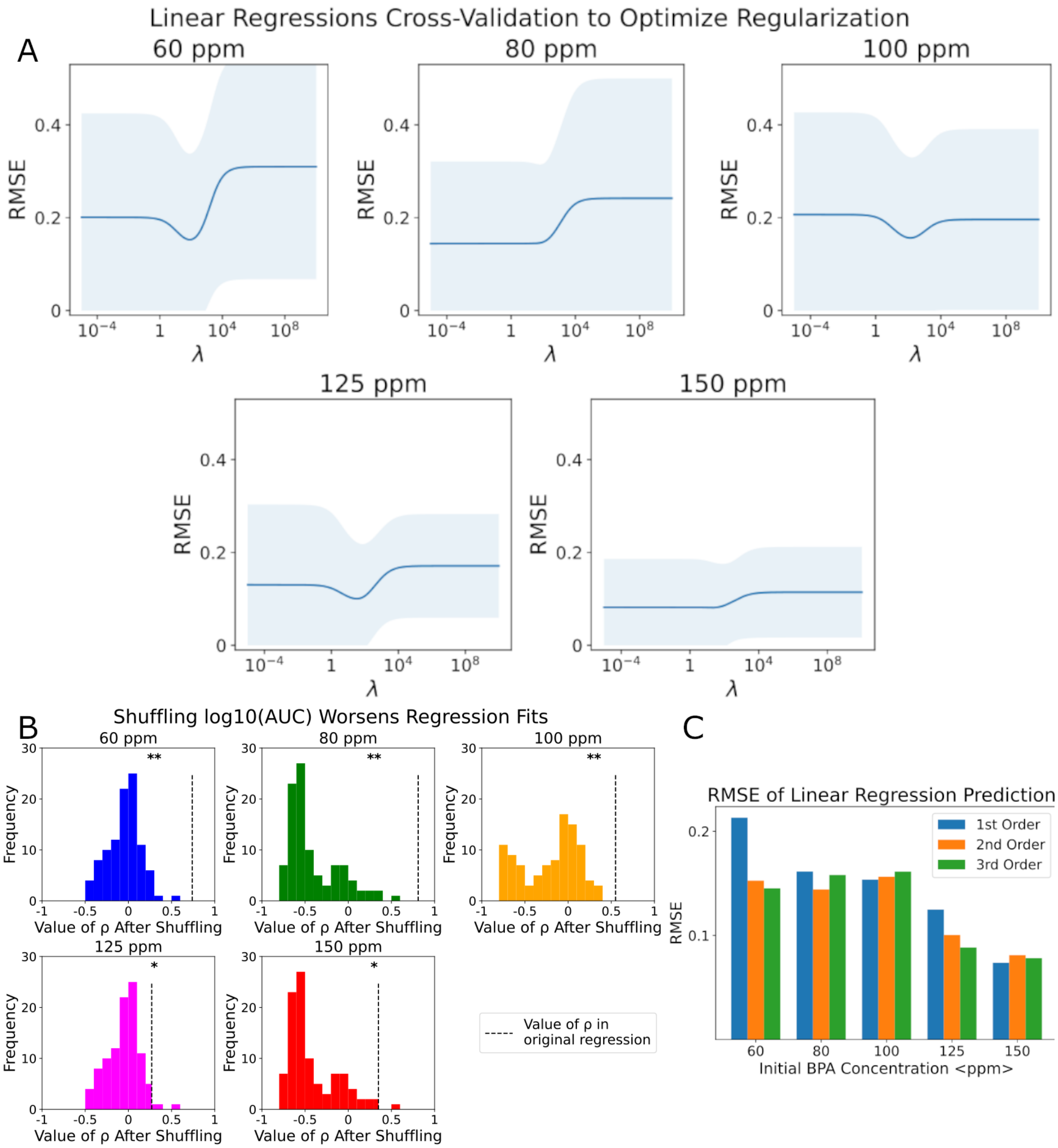


Figure S5: **Linear Regression Models.** See following page for caption.

Figure S5: (Continued from previous page) **(A)** Root Mean Squared Error (RMSE) (Eqn. 9) vs $\lambda$ during leave-one-out cross-validation for all five regressions (see Methods Tbl. 1, and Fig. 3). The blue line is the average across all folds, and the shaded region is the standard deviation across all folds. The optimal penalty term was selected for each model to minimize the RMSE. **(B)** Shuffling the response variable reduces the predictive power of the model. Each panel corresponds to different initial BPA concentrations. For each concentration, we shuffled the response variable (log10(AUC)) 100 times, for each shuffle we performed a regularized linear regression as described in the Methods. The histogram shows the distribution of Pearson correlation ($\rho$) between measured and predicted log10(AUC) for the 100 randomizations. Dotted black lines indicate the $\rho$ of the original regression fit (Fig. 3 B). *$p$-value $<$ .05, **$p$-value $<$ 0.01. Model fits to randomized data are significantly worse than the true data fits in Fig. 3. **(C)** Root Mean Squared Error (RMSE) for each of the five concentrations fit to linear regressions of models truncated at first-order terms ($\beta_i$), second-order terms ($\gamma_{i,j}$), and third-order terms ($\zeta_{i,j,k}$). For each model at each order, leave-one-out cross-validation was done to pick the optimal penalty term to minimize out-of-sample RMSE (see Methods).

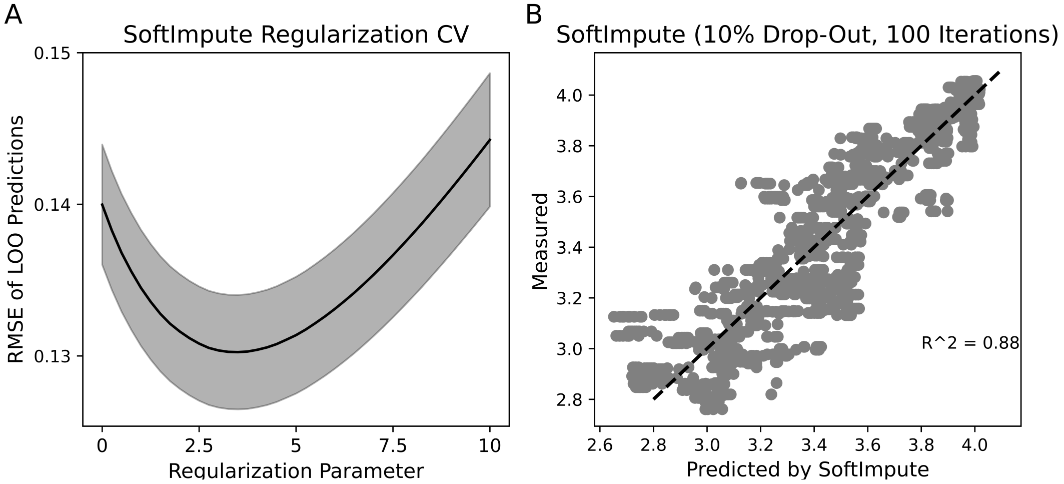


Figure S6: `SoftImpute` **Optimization** **(A)** Root Mean Squared Error of Leave-One-Out predictions as a function of the `SoftImpute` regularization parameter (see Methods). Regularization parameters were tested between 0 and 10 in increments of 0.25, and $R^2$ was calculated from the complete leave-one-out predictions of the AUC matrix. The optimal regularization of 3.5 was chosen for all imputations. Shaded region indicates the standard error of all folds. **(B)** `SoftImpute` predictions from randomly dropping 10% of community AUCs. Random drops were performed 100 times, and all resulting predictions are plotted. Dotted line represents y=x. $R^2$ = 0.88, $p$-value $<$ 0.01 as computed by the `scikit-learn` function `r2_score`.

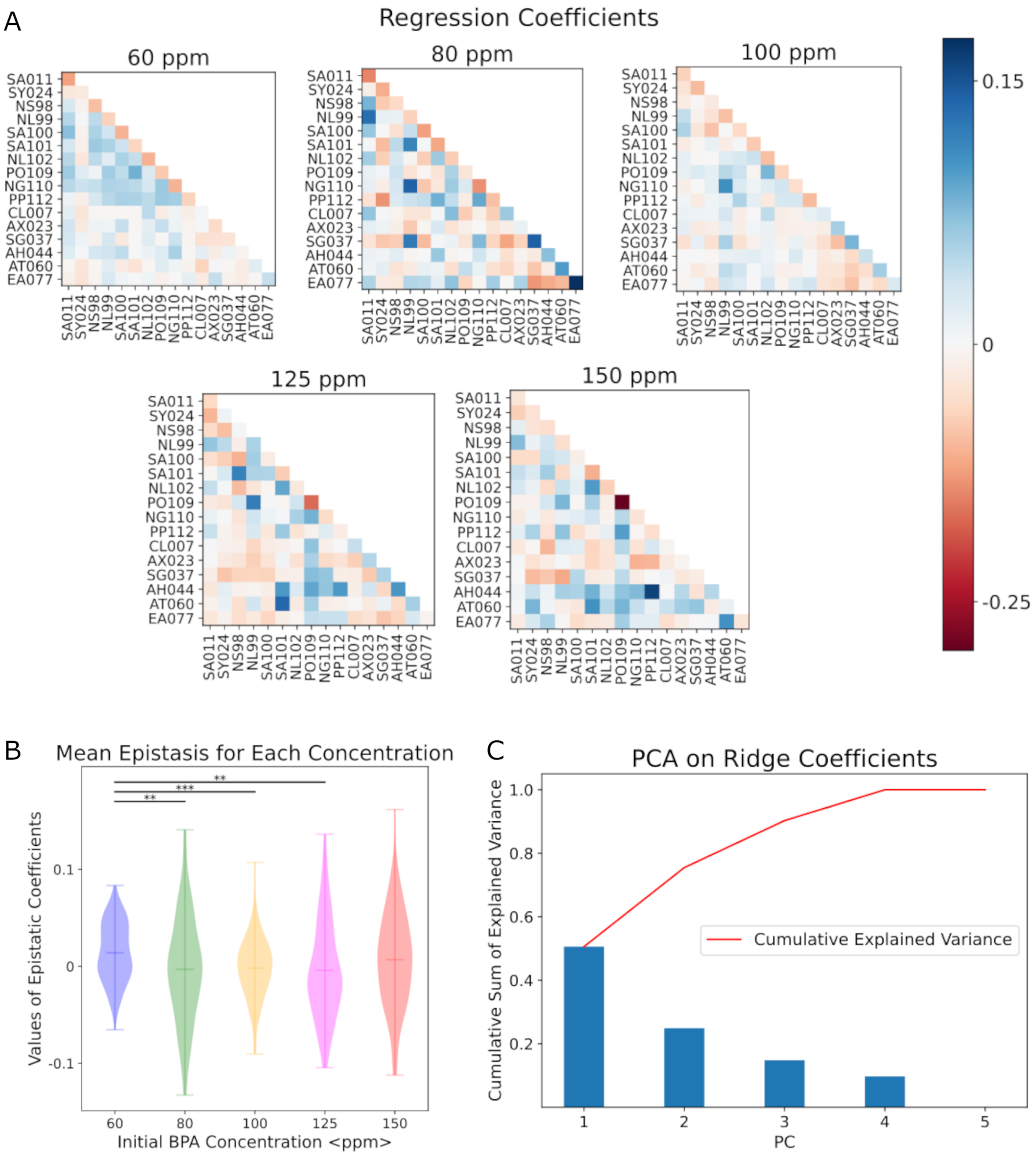


Figure S7: **Analyzing Regression Coefficients to Understand the Functional Landscape**

Figure S7: (Continued from previous page) **(A)** Heatmaps of regression coefficients from each of the five regression models (Fig. 3). Diagonal entries represent additive ($\beta_i$) coefficients, while off-diagonal entries represent epistatic ($\gamma_{i,j}$) coefficients. **(B)** Violin plots of epistatic coefficients ($\gamma_{i,j}$ terms) for each initial BPA concentration. More positive values indicate more antagonistic epistatic interactions (one strain inhibiting another), while more negative values indicate more cooperative epistasis (improving BPA degradation). Horizontal lines within distributions represent the mean coefficient. Stars indicate p-values for statistical tests of differences in average epistasis via two-sided t-test: $^{**}p < 0.01$, $^{***}p < 0.001$. At 150 ppm, the mean epistasis is not significantly different from 60 ppm, but the variance of the distribution of epistatic terms is larger (Fig. 4D). **(C)** PCA on Regression Coefficients show that the first two principal components explain 75% of the variation between coefficients of the five models, while the first four PCs explain all of the variation.

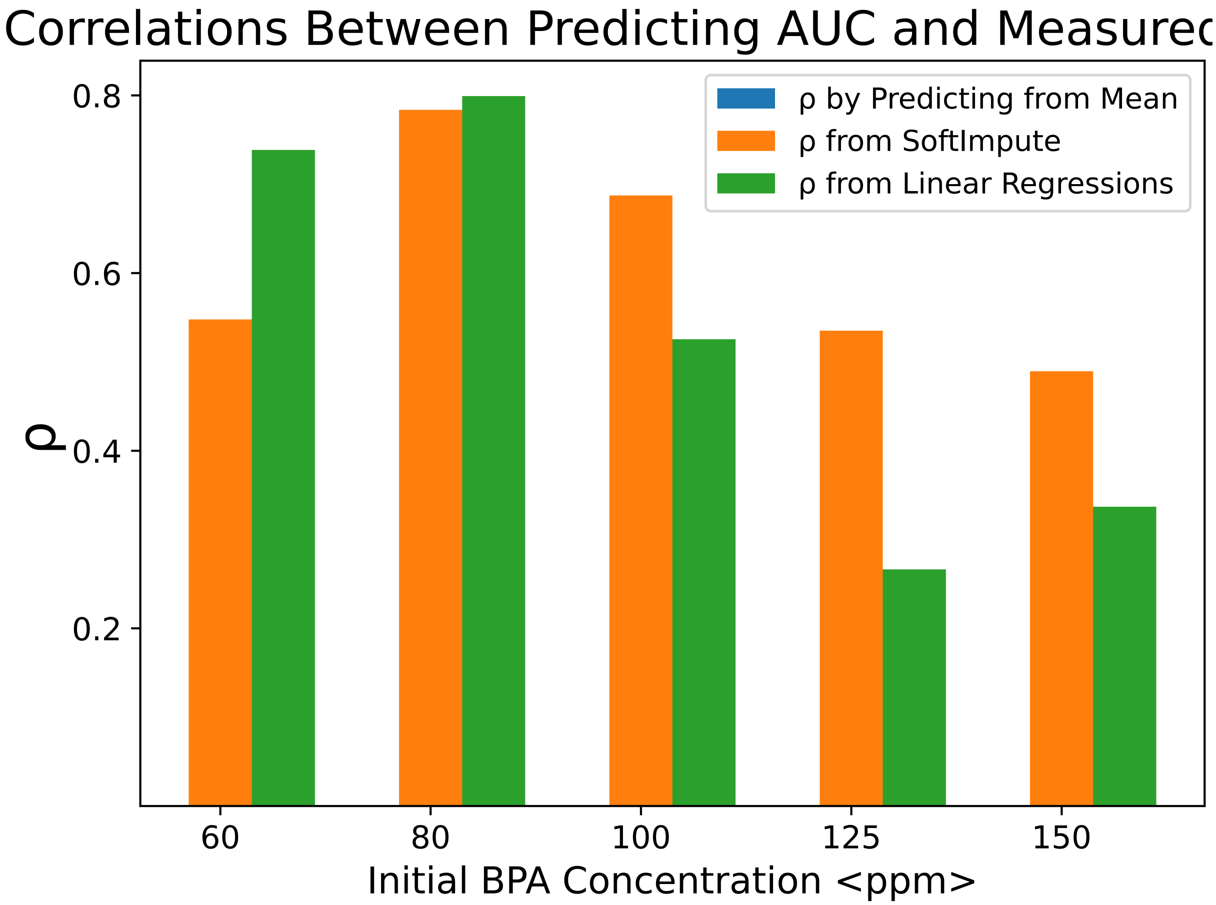


Figure S8: **Comparing Pearson's of Predicting by Mean, `SoftImpute`, and Regressions** Pearson's correlation between measured and predicted AUC for each of the five concentrations using three models: a model that predicts the average AUC for each initial concentration (blue), `SoftImpute` predictions (orange, Fig. 2 D), and regularized linear regressions (green, Fig. 3). Note that for the model predicting the average AUC, $\rho = 0$ for all five initial concentrations.

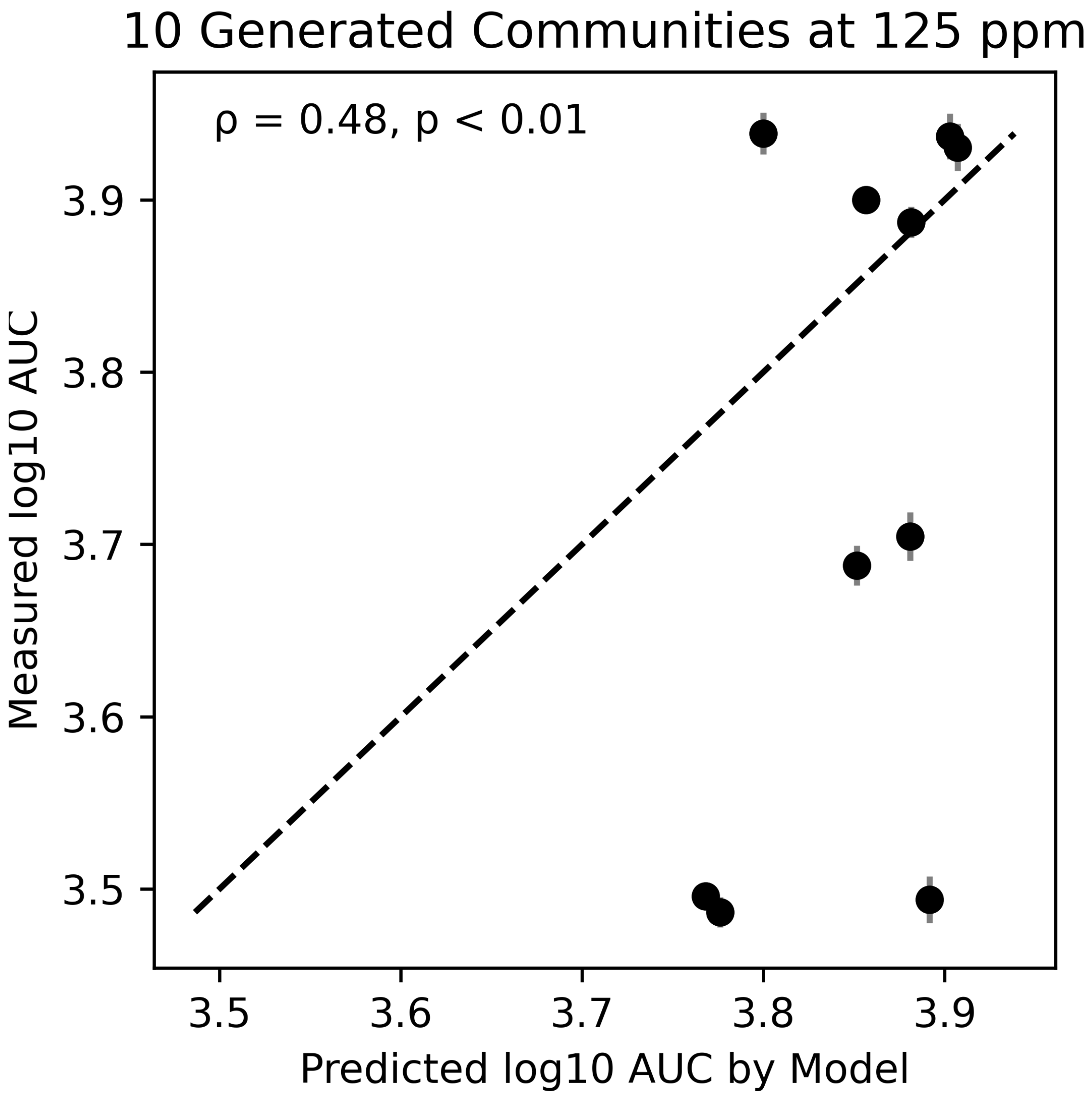


Figure S9: **Model Predicts Communities Designed at 125 ppm** AUC of BPA degradation in the same ten constructed communities as Fig. 3 at an initial concentration of 125 ppm BPA. Error bars represent the standard deviation of technical replicates. Dotted line shows $y = x$, perfect predictions. $p < 0.01$ from 100 bootstrapping iterations rejects the null hypothesis that the Pearson's correlation coefficient is $<0$.

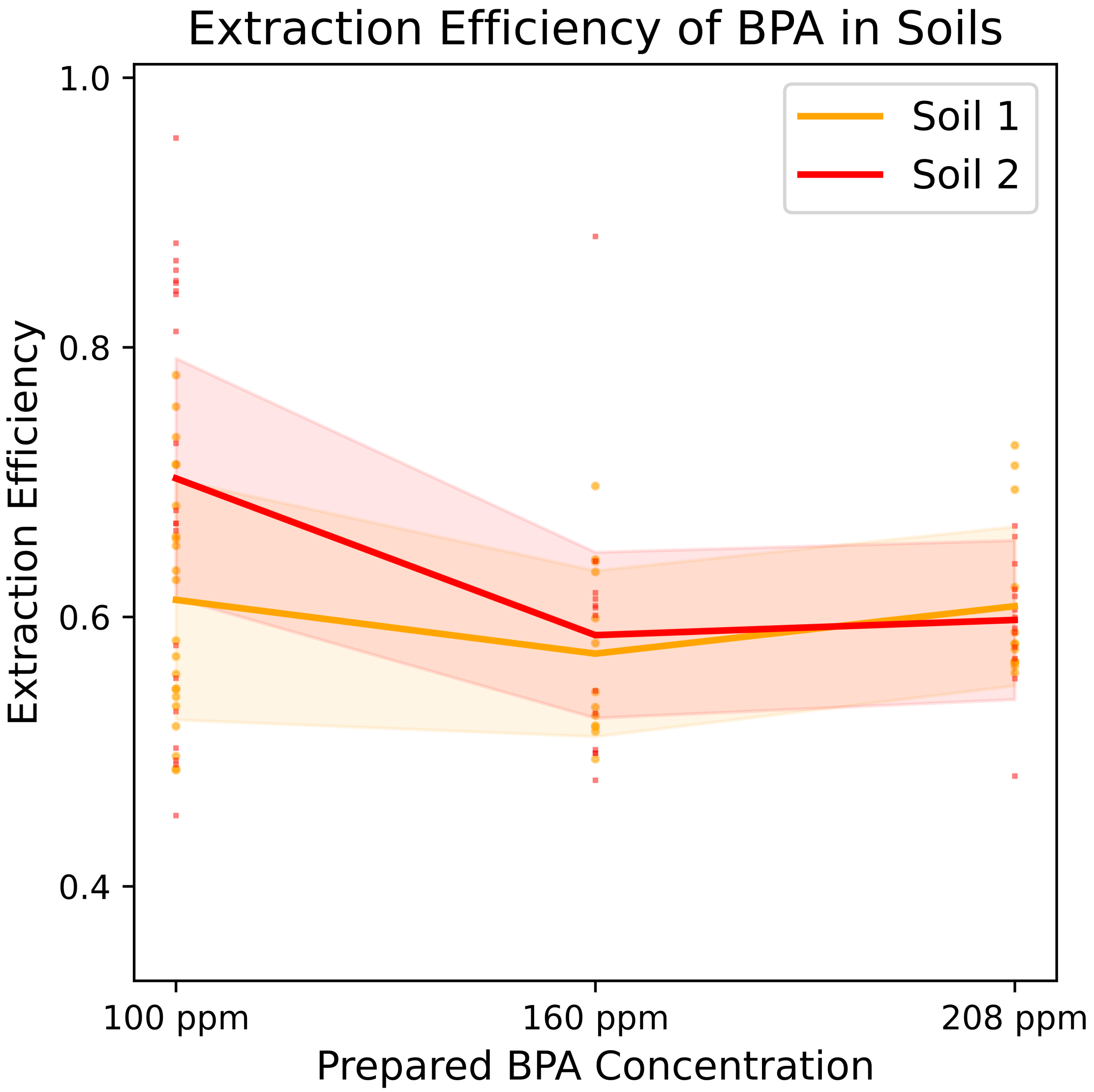


Figure S10: **BPA Extraction Efficiency from Soil Slurries** Extraction efficiency of BPA from both soil slurries from three initial BPA concentrations. Solid colored lines represent the mean extraction efficiency for each soil at each concentration, and shaded regions denote one standard deviation from the mean. Number of replicates varied between 20 and 40 for each condition. For extraction protocol see Methods.

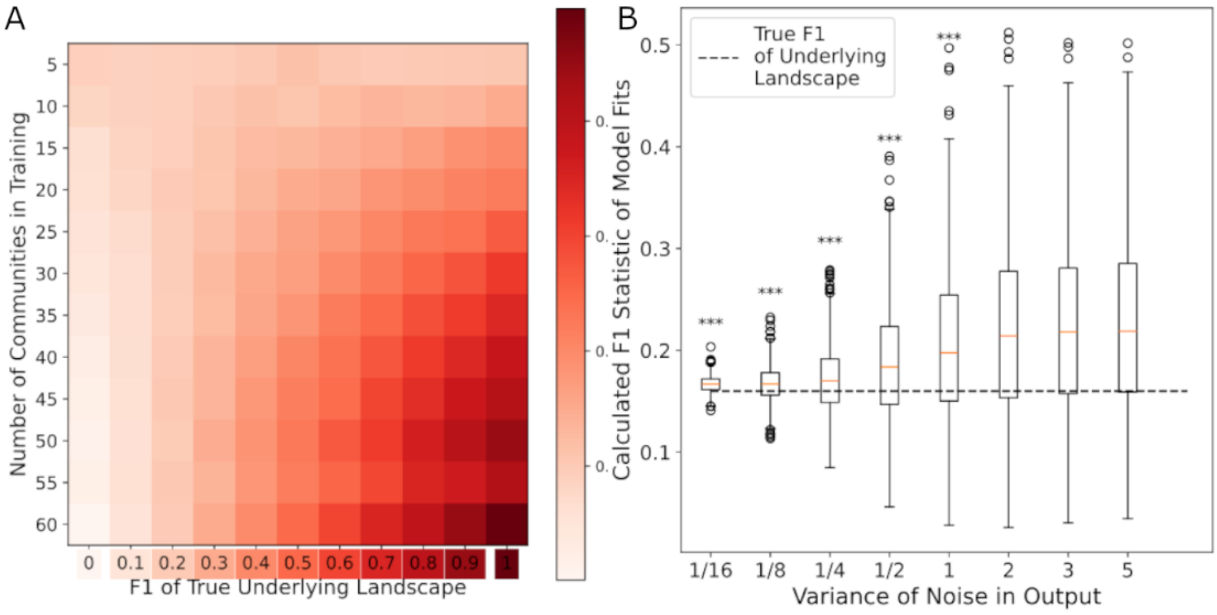


Figure S11: **Calculated $F_1$ Statistic Increases when Model Fits are Poor (A)** Heatmap showing the calculated $F_1$ statistic of regression models inferred from training sets of various sizes $s'$ and different imposed $F_1$ values (ruggedness), averaged across 100 random samplings. Darker colors indicate higher $F_1$ statistics (more additive and less epistatic). The resulting heatmap shows that $F_1$ can be accurately inferred as long as there are enough data points ($s'$) but quickly becomes inaccurate when there is too little data. For all entries, the error in the inferred $F_1$ is less than the mean. **(B)** Effect of poor model fits (represented by increasing $\sigma^2$ of the error term $\eta_c$) on the calculated $F_1$ statistics. Results from 1000 iterations show that for epistatic landscapes, increasing noise also increases the inferred $F_1$ statistic rather than decreases it. $^{***}p < 0.001$ decreased mean compared to $\sigma^2 = 5$ as determined by a one-sided t-test.

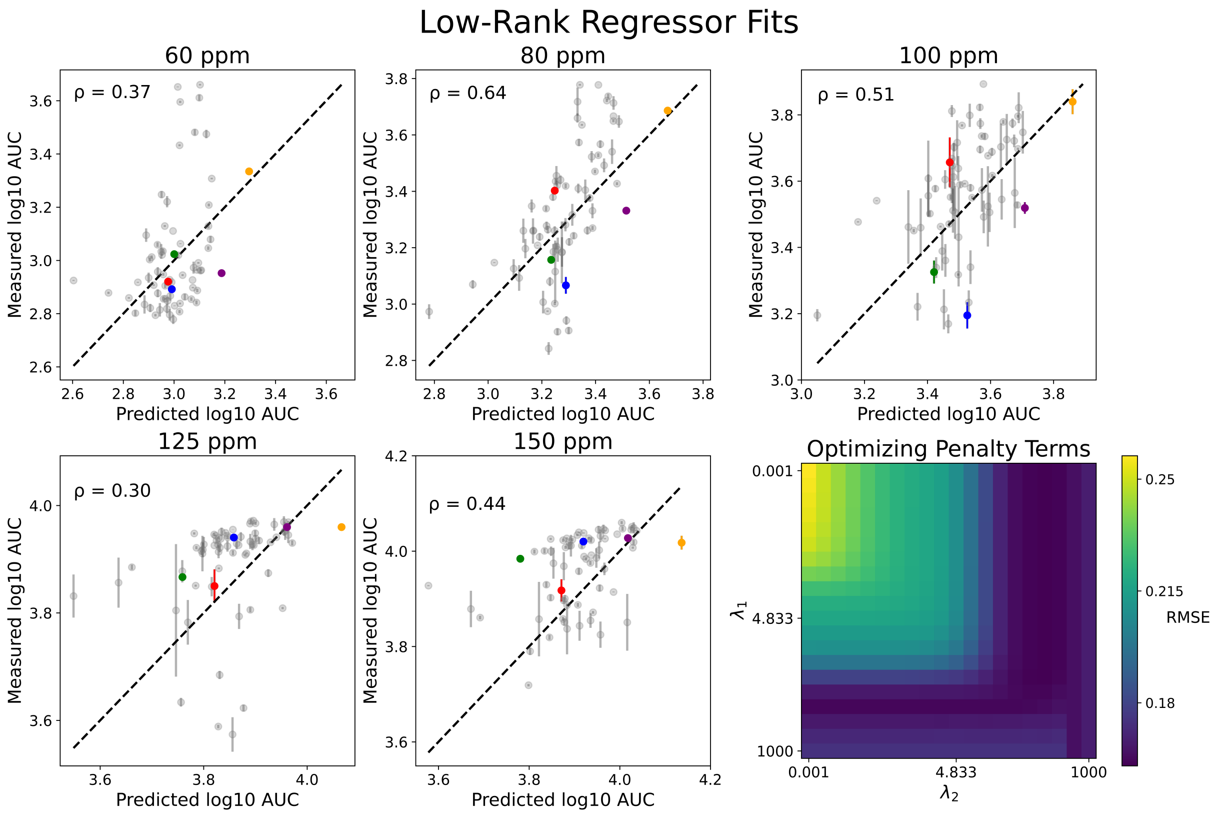


Figure S12: **Low-Rank Regressor Fits to BPA AUC Data** Low Rank Regressor fits to our AUC matrix (see Supplementary Information). *Top Row and Bottom Left and Center Panels* Fits for each of the five initial concentrations. Dotted black lines denote $y = x$ (perfect predictions) within each concentration. Vertical error bars represent the standard deviation of AUCs in technical replicates. *Bottom Right Panel* Total RMSE of all fits as a function of the regularization parameters $\lambda_1$ and $\lambda_2$ (Eqn. 9). Darker shades represent smaller RMSEs.
